# Real-Time Nanoscale Investigation of Spore Coat Assembly in *Bacillus subtilis*

**DOI:** 10.1101/2024.12.19.629407

**Authors:** Armand Lablaine, Dimitri Juillot, Ciarán Condon, Rut Carballido-López

## Abstract

Spores of *Bacillaceae*, ubiquitous in soil, can withstand extreme conditions virtually indefinitely. The spore coat, one of the most sophisticated multiprotein complexes built by bacteria, protects the spore from environmental stresses and predators. In *Bacillus subtilis*, assembly of the multilayered coat initiates at the forespore pole and is perceived as a stable process that progresses continuously until covering the entire forespore surface. In contrast, in *Bacillus cereus*, coat formation initiates in the midspore region and extends outward to the spore poles. Using Structured Illumination Microscopy, we monitored *B. subtilis* coat development in real-time at the single-sporangium level with lateral resolution of 70-nm. We found that late-synthesized proteins from the innermost coat layers first assemble in the midspore region and are subsequently displaced towards the poles. This process is coupled with a unique redistribution of pre-assembled coat material across the forespore surface and influenced by outer coat development, highlighting a dynamic interplay in coat layers co-construction.

## Introduction

To survive harsh conditions such as nutrient deprivation encountered during their life cycle, *Bacillaceae* species undergo a complex developmental process that results in the formation of spores - dormant, virtually immortal bacterial forms^1^. Sporulation is an irreversible and energetically costly survival strategy for a bacterium, involving an intricate sequence of cell differentiation events (Fig. 1A). First, the bacterial cell undergoes asymmetric division, resulting into a so-called sporangium composed of two compartments with unequal sizes and distinct fates: a small forespore (or prespore) that will mature into a resistant spore, and a larger mother cell that supports spore development (Fig. 1A, panel a). The mother cell then engulfs the forespore through a phagocytosis-like mechanism (Fig. 1A, panels b-c). Once engulfment is complete, the ovoid forespore is fully enclosed by the mother cell cytoplasm, isolating it from the external environment (Fig. 1A, panel d). After this step, the forespore undergoes progressive dehydration in preparation for dormancy, and develops refractile properties (Fig. 1A, panel e). Finally, the mother cell lyses releasing the spore into the environment (Fig. 1A, panel f). In the model Gram-positive bacterium *Bacillus subtilis*, the entire sequence of events typically spans 8 to 10 hours at 37°C, and is precisely regulated in time and space by the activation of alternative RNA polymerase sigma factors, which control differential gene expression in the two sporangial compartments. In the mother cell, SigE (σ^E^) is initially active and is replaced by SigK (σ^K^) upon completion of engulfment (Fig. 1A). Additional, ancillary transcription factors, such as GerR and GerE, fine-tune the timing of gene expression^2^. The stages of sporulation and the associated regulatory genes are largely conserved among *Bacillaceae* species^3,4^.

**Figure 1.**
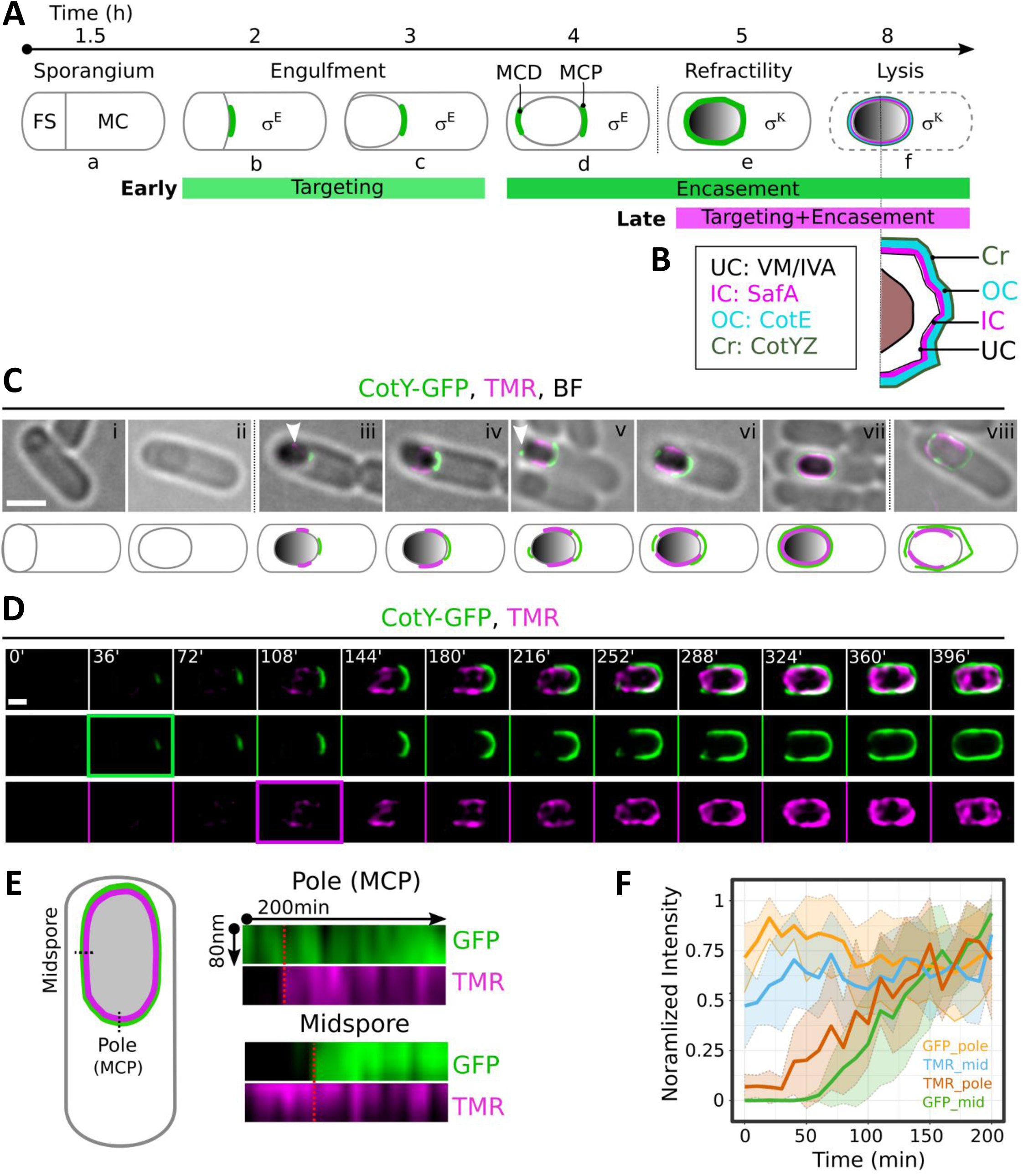
TMR-star and CotY-GFP initially target different region of the forespore surface. **(A)** Schematic of *B. subtilis* sporulation and coat development. First, the rod-shaped bacterium asymmetrically divides into a small forespore (FS) and a larger mother cell (MC), both defining a sporangium (a). Then, the MC engulfs the FS (b-d). Upon engulfment completion the FS is dehydrated and develops refractility (e). Ultimately, the MC lyses releasing the spore (f). In *B. subtilis*, the overall process takes 8 hours at 37°C. The MC-specific RNA polymerase sigma factors SigE (σ^E^) and SigK (σ^K^) support the production of coat proteins. Timing of targeting and encasement by early and late-synthesized coat proteins is represented by colored bars. **(B)** Schematic of a spore cross-section showing the different coat layers and associated morphogenetic factors: UC, under coat (black); IC, inner coat (magenta); OC, outer coat (cyan); Cr, crust (dark green). **(C)** Representative super-resolved localization patterns of CotY-GFP (strain PE3158, green) and TMR-star (magenta) in sporangia at various sporulation stages collected at hour 3 (panel i), 5.5 (panels ii to vi) and 7.5 (panels vii and viii) after resuspension in sporulation media. Note that BF illumination allows to identify the FS membrane prior refractility development (see Fig. S4B). Arrowheads, initial targeting of TMR-star on the midspore region and initiation of encasement by CotY-GFP. Scale bar, 1 µm. Schematics depicting the localization patterns of CotY-GFP and TMR-star signals are below each micrograph. **(D)** Single-sporangium time-lapse of CotY-GFP and TMR-star localization on an agarose-pad. A typical montage is shown with 36 min interval. The montage is representative of 3 independent experiments. The green boxed and magenta boxed frames indicate the first time point of CotY-GFP and TMR-star signal detection, respectively. The separate channels and complete sequence with original 12 min interval are shown in Fig. S2. Scale bar, 500 nm. **(E)** Representative kymograph analysis of the CotY-GFP and TMR-star signals over an 80 nm wide line perpendicular to the forespore surface, at either the center of the MCP pole or in the midspore region as depicted in the cartoon on the left. Kymographs were done over 200 min time-lapses. The first frame corresponds to the first signal detected (CotY- GFP at the MCP pole and TMR at the midspore). The red dashed lines indicate the delayed detection of the signal in the other channel (TMR at the MCP pole and CotY-GFP at the midspore region. **(F)** Intensity profiles of TMR-star and GFP signals at the points on the MCP pole and the midspore region shown in **(E)** over the 200 min time-lapses. Lines show the mean of 6 cells analyzed, and the shadowed area represents the SD.

Prior to its programmed death, the mother cell supports the formation of two protective structures around the forespore: the internal cortex, composed of modified peptidoglycan, and the extracellular proteinaceous coat. Strikingly, despite substantial conservation of genes encoding coat proteins, different coat structures have been described among *Bacillaceae* species, likely reflecting adaptation to diverse ecological niches^5,6^. The best-studied example is the multilayered coat that shields the surface of *B. subtilis* spores (Fig. 1B). This structure, approximately 200 nm thick, is built through contributions of at least 80 proteins (with ∼15% of sporulation genes coding for coat components^5^), rendering the coat one of the most complex protein assemblies in bacteria. In contrast, species within the *Bacillus cereus* group possess a single coat layer encased by a loose-fitting balloon-like exosporium. Together, the proteinaceous surface layers form an extracellular shell that acts as a semi-permeable barrier, controlling the transit of molecules to prevent germination in suboptimal environments, which would threaten the survival of the future colony^7,8^. These layers also confer resistance against digestion by protozoans, nematodes and macrophages^9–11^. Additionally, they modify the hydrophobicity properties of the spores, which are important for their environmental dissemination^12–14^.

Based on genetic and electron microscopy analysis, the *B. subtilis* spore coat has been divided into four distinct layers, from the inner to the outermost: i) the undercoat (or basement layer), which was recently visualized for the first time using advanced cryo-electron tomography^15^, ii) a lamellar inner coat, iii) a highly electron-dense outer coat, and iv) a glycoproteic crust^5,16^ (Fig. 1B). The assembly of each layer relies on specific morphogenetic proteins: SpoIVA and SpoVM drive assembly of the undercoat, SafA of the inner coat, CotE of the outer coat, and CotYZ are required for crust formation^5,16,17^ (Fig. 1B). Assembly of the inner and outer coat layers appears largely independent, as deletion of *safA* has minimal impact on outer coat formation, and deletion of *cotE* does not affect inner coat assembly^18,19^. The mechanism of coat assembly in *B. subtilis* was extensively studied using conventional epifluorescence microscopy and a collection of over 50 GFP-tagged coat proteins that were visualized in different mutants of the pathway^5,16,20–23^. These studies suggested that coat protein assembly begins at the spore pole, primarily driven by its particular positive curvature, and spreads from this initial location supported by self-assembly and successive transcription waves^5,16,21,23–26^. Initially, at the “targeting step”, early-synthesized (σ^E^-dependent) coat proteins, which include the morphogenetic proteins of all four coat layers, assemble into a polar cap at the mother cell proximal pole (MCP) of the forespore^21,23,27^ (Fig. 1A, panels b and c, green cap). After engulfment is completed, most early-synthesized proteins form a second cap on the newly formed mother cell distal (MCD) pole of the forespore (Fig. 1A, panel d), and then spread from the two forespore poles towards the midspore region (Fig. 1A, panel e). This progression from one polar cap to a complete shell is referred to as “encasement”^21^. Unlike the early-synthesized coat proteins, the assembly of late-synthesized (σ^k^-dependent) coat proteins is particularly difficult to resolve in conventional fluorescence microscopy images and remains unclear. This is also true for some early-synthesized inner coat proteins such as YutH, YsxE or YisY^23,28^. In *B. cereus* species, orthologues of the *B. subtilis* morphogenetic coat proteins are present and play critical roles in coat and exosporium formation^29–37^. However, unlike *B. subtilis*, where coat assembly begins at the forespore pole, in *B. cereus* species, coat material first appears in the midspore region^37–39^, suggesting that coat formation follows different dynamics across *Bacillaceae* species.

Our current understanding of *B. subtilis* coat development suffers from several major limitations, particularly the resolution of conventional light microscopy and the inability to track protein dynamics throughout sporulation at the single-sporangium level. The diffraction limit of visible light restricts lateral resolution to approximately ∼200-250 nm, close to the ∼200 nm thickness of the *B. subtilis* coat, making it impossible to distinguish individual coat layers (inner coat, ∼40-70 nm; outer coat, ∼70-200 nm^40^) using conventional fluorescence microscopy. Sub-diffraction localization of individual coat proteins has only been reported in sporangia at a late sporulation stage or in mature spores via computational image analysis methods. Furthermore, while early and intermediate sporulation stages can be monitored using commercial membrane dyes and phase-contrast microscopy; respectively, no physiological markers are currently available to monitor later stages once the forespore becomes refractile. Yet this refractile stage, which lasts about 3 hours (Fig. 1A, panels e-f), is by far the longest stage of sporulation and poses significant challenges due to the risk of premature germination^41^. Additionally, mid-term (hours-long) single-cell time-lapse imaging, which is the time-scale of coat development, has not yet been applied to investigate coat assembly. Because of these limitations, coat proteins exhibiting dynamic behavior on the forespore surface during sporulation would have gone undetected.

In this study, we performed single-cell time-lapse analysis of coat development during *B. subtilis* sporulation using Structured Illumination Microscopy (SIM), which doubles the spatial resolution of conventional epifluorescence microscopy (SIM theoretical lateral resolution, ∼120 nm^42^). Furthermore, we employed an advanced lattice illumination pattern (lattice-SIM, see Methods), which reduces phototoxicity and, combined with the dual iterative SIM² deconvolution algorithm, doubles the standard SIM lateral resolution^43^. Using this approach, we successfully imaged the entire coat development process at the single-cell level over more than 10 hours with a measured lateral resolution of ∼70 nm. We also demonstrated that TMR-star, a cell-permeable red fluorescent ligand typically used to label the SNAP-tag, which was recently shown to bind to the coat of *B. cereus* species^35,39^, specifically labels the inner coat of *B. subtilis*. Notably, the TMR-star signal was initially detected in the midspore region and eventually covered the entire forespore surface, similar to *B. cereus* coat deposition dynamics^33,35,37,38^. Using 2D and 3D dual-color lattice SIM², we then visualized a collection of GFP-tagged coat proteins from all sublayers relative to TMR-star-labelled inner coat deposition, and found that late-synthesized proteins from the inner coat layer are also initially detected in the midspore region and colocalize with TMR-star throughout sporulation. Furthermore, the initial detection TMR-star in the midspore region was synchronized with an unexpected redistribution of early-synthesized inner coat proteins, constituting the first observation of apparent protein relocalization on the forespore surface. Finally, we show that the outer coat assemby influences the localization dynamics of early-synthesized inner coat proteins. Altogether, our findings redefine and unify the models of coat formation among *Bacillaceae* species.

## Results

### TMR-star as a marker of late sporulation sub-stages

*B. subtilis* coat assembly has been extensively studied at the population level, by imaging sporulating cells collected at different time points during sporulation. Using commercial membrane fluorescent dyes and phase contrast microscopy, it was possible to determine a virtual sequence of assembly for various GFP-tagged coat proteins^5,16,29^. However, for many coat proteins, particularly those assembling after the forespore becomes refractile (late-synthesized proteins), membrane dyes and phase contrast do not permit identification of intermediate assembly stages. To overcome this limitation, we sought a new physiological marker that could reliably distinguishing later stages of *B. subtilis* sporulation and allow for continuous live-cell SIM imaging. We recently showed that TMR-star, a synthetic red fluorescent ligand originally developed to label SNAP-tagged proteins, binds to the coat of *B. cereus* and *Bacillus thuringiensis* once the forespore becomes refractile^39^. Since the *B. subtilis* coat has been reported to bind various fluorescent dyes^41,44^, we tested TMR-star labelling in sporulating *B. subtilis* cells collected at various times after resuspension in sporulation medium, and imaged their midplane section using 2D-Lattice-SIM². As a control, we imaged a functional GFP fusion to the morphogenetic protein CotY, previously shown to localize to the crust layer^17^, expressed ectopically from the *amyE* locus under control of its native promoter. CotY-GFP assembly followed the expected sequence for early-synthesized coat proteins, first forming a cap at the MCP pole, then as a second smaller cap at the MCD pole, and eventually a complete ring around the spore mid-section (Fig. 1C). The CotY-GFP MCP cap assembled late, only appearing in sporangia with a refractile forespore (also referred to as refractile sporangia), unlike other morphogenetic coat proteins that localize as a polar cap before engulfment completion^23,27^. Similar to CotY-GFP, the TMR-star signal was detected only in refractile sporangia (Fig. 1C, panels iii-vii). However, TMR-star first appeared in the midspore region (arrowhead in panel iii), and not at the MCP pole like CotY-GFP, before spreading bidirectionally along the long axis of the forespore (Fig. 1C, panels iv-vi). When both CotY-GFP and TMR-star signals fully encased the forespore, the TMR-star signal was clearly observed beneath the CotY-GFP signal, with no apparent overlap between the two (Fig. 1C, panel vii). In sporangia undergoing premature germination, both CotY-GFP and TMR-star signals were discontinuous and disorganized (Fig. 1C, panel viii). This analysis reveals an original TMR-star localization pattern, binding to an unidentified substrate under the spore outermost layer. Notably, the dynamics of TMR-star localization and its internal positioning relative to the glycoproteinaceous crust, strongly resemble the sequence of coat assembly described in *B. cereus* species, where the TMR-star signal lies beneath the exosporium layer^35,37,39^. Since TMR-star allowed us to distinguish several substages in refractile *B. subtilis* sporangia, we concluded that it can be used as a physiological marker for the late stages of *B. subtilis* sporulation.

### Analysis of *B. subtilis* coat development using single-cell time-lapse dual color Lattice-SIM^2^

Mid-term single-cell time-lapse microscopy has been successfully used to examine bacterial cell communication and memory during sporulation^45–48^. However, it has not yet been used to monitor spore coat development, nor has it been combined with super-resolution microscopy. To capture the dynamics of CotY and TMR-star on the spore surface more precisely, we monitored the CotY-GFP and TMR-star signals at the single-sporangium level over extended periods (up to 10 hours) with a temporal resolution of 12 minutes. Leveraging the particular robustness and reproducibility of sporulation development in microcolonies grown on agarose-coated microscopy slides (Fig S1A), which significantly reduced premature germination events compared to sporangia collected from liquid cultures (Fig S1B), we conducted minimally invasive lattice-SIM imaging (see Methods).

The time-lapse sequence of CotY-GFP and TMR-star assembly events in cells immobilized on agarose pads (Fig. 1D, see also Fig. S2 for the individual channel sequences) mirrored the sequence derived from cells collected from liquid cultures (Fig. 1C). CotY-GFP was initially detected at the MCP pole, while TMR-star first localized at the midspore region, and both signals then spread from these initial sites and stably assembled over the entire forespore surface (Fig. 1D-F and Fig. S2). We detected CotY-GFP earlier (green square at 36 min in Fig. 1D) than TMR-star (magenta square at 108 min in Fig. 1D), while TMR-star completed spore encasement before CotY-GFP (Fig. S2). From their initial detection on the forespore surface, CotY-GFP and TMR-star required 255±42 min and 138±20 min (mean±SD; n=22), respectively, to fully encase the forespore under these conditions (see Methods). We noticed significant variability in the timing of CotY-GFP MCD cap formation, which occurred 115±43 min after MCP cap formation. Taken together, these results demonstrate that lattice-SIM² enables single-cell monitoring of spore coat formation and that TMR-star can serve as an effective physiological dye for super-resolved time-lapse imaging of late sporulation stages. Additionally, they show that assembly of the coat layer to which TMR-star binds initiates in the mid-forespore region and occurs faster than crust assembly.

### TMR-star stains the inner coat of *B. subtilis* spores

In *B. cereus* species, TMR-star was shown to specifically localize to the single spore coat layer^35,39^. Here, we observed that, in *B. subtilis,* TMR-star binds to a layer of the refractile forespore located beneath the CotY-GFP crust signal (Fig. 1C, panel vii). To determine whether TMR-star labels a specific layer or component of the multilayered *B. subtilis* coat, we performed dual-color nanoscale analysis using GFP fusions to proteins of the four distinct coat layers (Fig. S3A)^5^: the undercoat (YhaX), the inner coat (YmaG), the outer coat (SpsI), and the crust (CotY). For this analysis, we focused on sporangia collected 6-7 hours after resuspension, in which TMR-star labelled the entire forespore periphery in midplane sections. To further refine the spatial mapping of these markers, we imaged sporangia immobilized vertically in agarose microholes, to better appreciate the relative position of the different markers around the midspore circumference (Fig. 2A).

**Figure 2.**
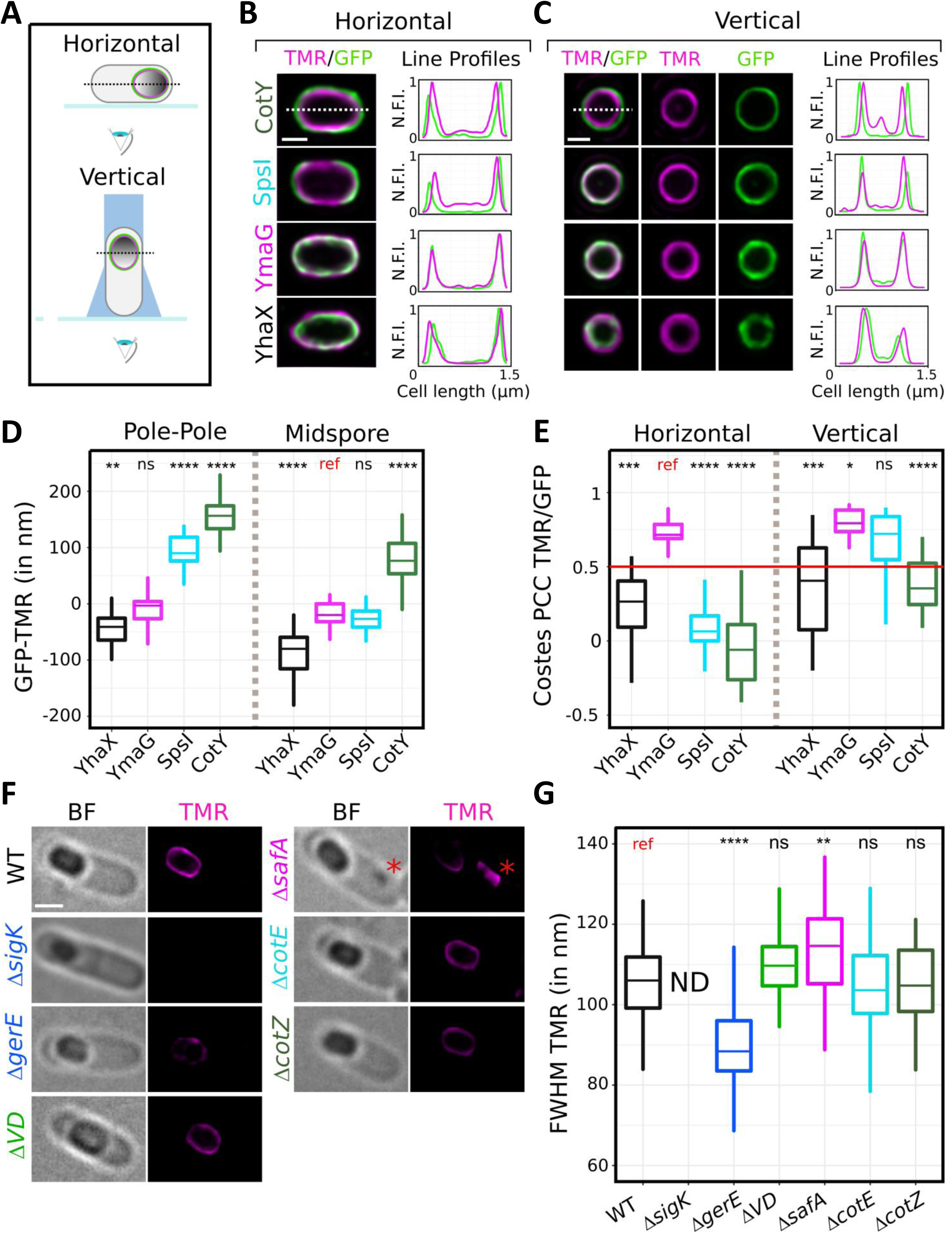
TMR-star stains the inner coat of B. subtilis spores. **(A)** Schematic representation of the experimental setup used to image horizontally (top) or vertically (bottom) immobilized sporangia using agarose nanoholes. The medial focal plane selected for imaging is represented by a dotted line, and the coverslip as a thick blue line. **(B, C)** *Left panels*, representative super-resolution micrographs of sporangia cross-sections completely encased by TMR-star signal (in magenta) and GFP fused to the indicated coat proteins (in green, CotY, PE3158; SpsI, PE2708; YmaG, CF55; YhaX, PE388) lying horizontally (B) or vertically immobilized (C). Scale bar, 500 nm. *Right panels*, fluorescence intensity line profiles (dashed white line) of TMR-star and GFP signals. N.F.I, normalized fluorescence intensity relative to the maxima. **(D)** Distribution of the distance (in nm) between the GFP signal (fused to the indicated coat protein) and the TMR-star signal at the poles and at the midspore region of horizontally immobilized cells. At least 10 sporangia were analyzed per condition. **(E)** Distribution of Costes Pearson Correlation Coefficient (Costes PCC) between signals of TMR-star and GFP fused to the indicated coat proteins. At least 22 and 16 sporangia per strain were analyzed for horizontally and vertically immobilized cells, respectively. PCC=1 would indicate perfect colocalization; PCC=0, no colocalization; PCC>0.5 is considered as a significant colocalization (red line). **(F)** Micrographs of sporangia representative from the indicated genetic background (WT, PE479; Δ*sigK*, RCL1720; Δ*gerE*, RCL1696; Δ*spoVD* (Δ*VD*), RCL1697; Δ*safA*, RCL1698; Δ*cotE*, RCL1699; Δ*cotZ*, RCL1700) and labelled with TMR-star. A red asterisk indicates TMR-star signal abnormally localizing in the MC cytoplasm. BF, brightfield. Scale bar, 500 nm. **(G)** Distribution of Full Width at Half Maximum (FWHM) of TMR-star signal measured on the forespore surface in the indicated genetic background. ND, not detectable. At least 30 sporangia were analyzed per condition. **(B-G)** Sporangia from at least 2 independent experiments were collected between hours 6.5 and 7 after resuspension in sporulation medium. The non-parametric Mann–Whitney U test was used with the condition indicated as a reference (ref); ns: not significant; *P˂0.05; **P˂0.005; ****P˂0.00005.

As a control, we first checked whether coat shape and thickness were affected by the expression of the GFP-tagged coat proteins (Fig. S3B and C). Among the GFP-coat fusions tested, only YhaX-GFP expression affected forespore morphology, resulting in slightly longer forespores (Fig. S3B). We then assessed the lateral resolution of our lattice-SIM² set-up using a GFP-tagged single transmembrane domain (Nrny-sfGFP, see Material and Methods), and found it to be approximately 70 nm (Fig. S4). The Full Width at Half Maximum (FWHM) of the individual GFP fusions signals was ∼90-100 nm, with the exception of YhaX-GFP that we resolved down to ∼80 nm (Fig. S3D). These measures were all above the lateral resolution of our system, suggesting that the individual coat proteins do not polymerize as a single monolayer but rather organize into a network within their respective coat sublayers. We next examined the co-localization of GFP and TMR-star signals in both horizontally positioned cells on agarose pads and vertically immobilized cells in microholes (Fig. 2A-C). We measured the distance between the TMR-star and GFP signals at the forespore poles and midspore, respectively, (Fig. 2D) and computed their colocalization by measuring Costes Pearson correlation coefficient (PCC)^49,50^ for the fluorescence signals in the two channels (Fig. 2E). As visually observed in Fig 1C, panel vii, the crust reporter fusion CotY-GFP was positioned outside the TMR-star signal (Fig. 2B-D, see line profiles). Similarly, the outer coat reporter SpsI-GFP localized externally relative to the TMR-star signal at the forespore poles (Fig. 2B and D), but appeared largely superimposed with TMR-star in the midspore region (Fig. 2C and D). This is consistent with earlier TEM reports suggesting that the outer coat is thinner in this region^40^. In contrast, signals from the inner coat reporter YmaG- GFP and TMR-star clearly overlapped in both vertically and horizontally positioned sporangia (Fig. 2B-D). Finally, the undercoat reporter YhaX-GFP was localized internally to TMR-star in both cell orientations (Fig. 2B-D). Costes PCC confirmed co-localization between TMR-star and the inner coat protein YmaG (as well as SpsI in vertically oriented cells) (Fig. 2E). The same results were observed for other inner coat proteins tested such as YeeK and CotD (Fig. S5), while minimal colocalization was observed with CotY, YhaX, and SpsI, especially at the cell poles (Fig. 2E). Collectively, these data confirm that lattice-SIM² allows resolution of the distinct layers within the *B. subtilis* coat and show that TMR-star binds a substrate localized to the inner coat. Furthermore, imaging both horizontally and vertically immobilized sporangia provided further evidence that the outer coat is thinner along the forespore sidewalls compared to the poles.

### Genetic dependencies of TMR-star localization reflect those of a late-synthesized inner coat protein

We next investigated the genetic dependencies of TMR-star localization in *B. subtilis* sporangia. To this end, we analyzed TMR-star localization in mutants lacking (i) the late mother cell-specific transcription factors SigK and GerE, (ii) the major coat morphogenetic proteins SafA, CotE and CotZ, and (iii) the cortex formation determinant SpoVD. We focused on late transcription factors since TMR-star signals were detected only in refractile forespores (Fig. 1C). TMR-star localization was fully dependent on SigK and partially dependent on GerE (Fig. 2F and G). In the *ΔgerE* mutant, the TMR-star signal on the forespore periphery was thinner relative to the wild-type background (Fig. 2F and G, 88±11 nm, n=72 and 107±11 nm, n=46, respectively), suggesting an abnormal, thinner coat layer to which TMR-star binds. In the *ΔsafA* mutant, TMR-star partially covered the forespore surface and additionally appeared as non-adherent material in the mother cell cytoplasm (Fig. 2F, red asterisk). The TMR-star signal around the forespore was also slightly wider (Fig. 2G, 115±13 nm, n=41), suggesting abnormal organization of the structure to which TMR-star binds. In contrast, in *ΔcotE, ΔcotZ*, and *ΔspoVD* mutant sporangia, TMR-star signal localization and the thickness were comparable to wild-type cells (Fig. 2F and G). We concluded that the assembly of the substrate of TMR-star in the inner coat depends on late mother cell transcription factors.

### Assembly of late-synthesized inner coat proteins, like TMR-star staining, initiates in the midspore region

The genetic dependence and initial detection of TMR-star in the midspore region prompted us to hypothesize that some late-synthesized inner coat proteins might display the same localization dynamics than TMR-star on the spore surface. To test this, we performed single-cell time-lapse imaging of GFP fusions to the late-synthesized inner coat proteins YmaG, LipC and YeeK, which, like the TMR-star substrate, are σ^K^- and GerE-dependent^5^. We completed this analysis by imaging cells collected between hours 5.5 and 7 after resuspension in sporulation media and showing TMR-star signal only in the midspore region (Fig. S6A).

Expression of these GFP fusions did not affect the forespore morphology or the inner coat thickness (Fig. S3B and C) and TMR-star staining did not alter the localization of GFP-fusions (Fig. S6B).

YmaG-GFP was first detected in the midspore region, like TMR-star (Fig. 3A, B, see also Fig. S7 for individual channels with the original temporal resolution). On average, the YmaG-GFP signal appeared 44±20 min (n=11) after the initial detection of TMR-star in this region (Fig. 3B, midspore, the red dotted line indicates the initial detection of TMR-star, see also Fig. S7). YmaG-GFP signal next expanded bidirectionally toward the forespore poles alongside with TMR-star (Fig. 3A and B). Under our experimental conditions, YmaG-GFP fully covered the forespore surface within 135±35 min (n=21). LipC-GFP displayed a similar localization pattern, and completely covered the forespore within 215 ±25 min (n=16) following initial detection at the midspore (Fig. 3A and B, see also Fig. S7). Finally, while a weak YeeK-GFP signal initially covered the forespore surface uniformly, a more intense signal was detected on the forespore long axis concomitant with the TMR-star signal and spread bidirectionally with it (Fig. 3A and B, see also Fig. S7, outlined white areas), completely encasing the forespore within 181±27 min (n=16). In line with these observations, a convincing and continuous colocalization (Costes PCC>0,5) between GFP-tagged YmaG, LipC and YeeK and TMR-star was recorded on the forespore surface throughout 200 minutes after TMR-star detection in single-cell time-lapse movies (Fig. 3C). Overall, these findings reveal that both TMR-star and late-synthesized inner coat proteins initially localize to the midspore and share similar encasement dynamics. Further supporting this, strong colocalization between YmaG, LipC and YeeK with TMR-star was also observed in sporangia collected 5.5-7 hours after resuspension (Fig. S6C). This latter analysis also showed that the late-synthesized inner coat proteins YxeE and, to a lesser extent, CotD colocalize with TMR-star signals at an intermediate stage of its assembly, while, surprisingly, the early-synthesized (σ^E^-dependent) inner coat proteins YutH, YsxE and YisY did not (Fig. S6A and C).

**Figure 3.**
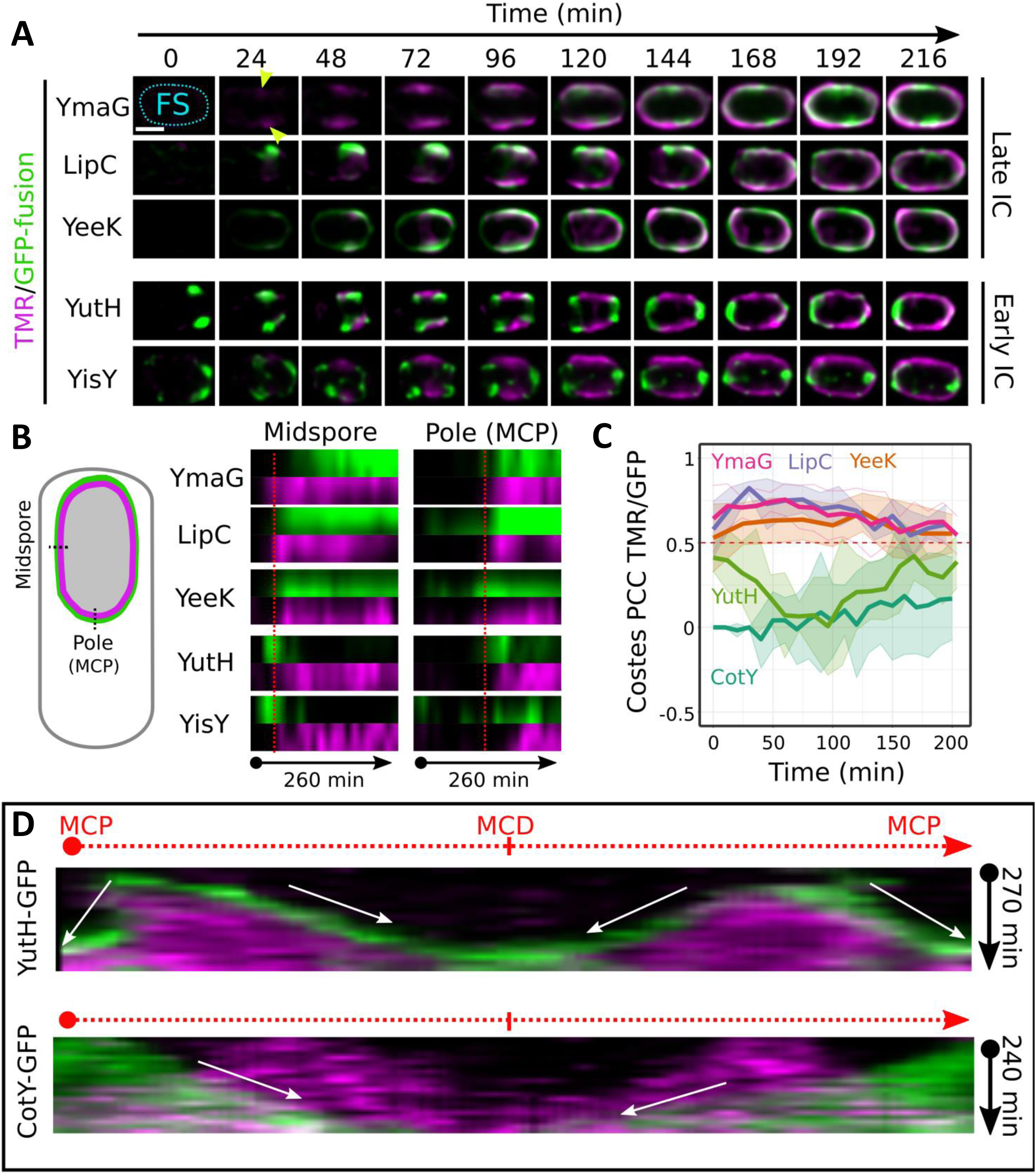
Assembly dynamics of inner coat proteins. **(A)** Single-sporangium time-lapses of TMR-star (magenta) and GFP (green) fused to late-synthesized (Late IC) inner coat proteins YmaG (CF55), LipC (PE1146), and YeeK (PE789) or early-synthesized (early IC) YutH (PE479) and YisY (PE485) during sporulation with 24 min intervals. See Fig. S7 and S8 for individual channels with original time resolution. Scale bar, 500nm. **(B)** Kymograph analysis of the GFP fusion to the indicated inner coat protein and TMR-star. Signals were measured along an 80 nm line perpendicular to the forespore surface, at either the center of the MCP pole or in the midspore region as depicted in the cartoon over 260 min in the time-lapses shown in (A), with time 0 min (one frame before TMR-star signal detection) as an initial point. The red dashed lines indicate the timing of detection of the TMR-star signal. **(C)** Evolution of Costes Pearson Correlation Coefficient (PCC) between super-resolved signals of TMR-star and GFP fused to the indicated coat proteins over time in individual sporangia. The red dashed line indicates a score of 0.5; above this score, the colocalization is considered significant. The bold profile is the mean of at least 5 sporangia per condition; the shadow represents the SD. Time 0 represents the time of TMR-star detection in movies. **(D)** Kymograph analysis of the YutH-GFP/TMR-star (PE479) and the CotY-GFP/TMR-star (PE3158) pairs along the entire forespore perimeter over 270 min and 240 min, respectively, time-lapses. White arrows indicate the different migration fronts, used to compute the redistribution/assembly speed of the GFP fusions. The four fronts of YutH-GFP represent a bidirectional redistribution from the midspore region towards the two poles, while the two fronts of CotY-GFP represent a mainly unidirectional assembly process from the MCP pole to the MCD pole.

### Early-synthesized inner coat proteins YutH and YisY are dynamically redistributed on the forespore surface

Previous diffraction-limited studies indicated that YutH, its paralogue YsxE, and YisY initially localize at the forespore pole, but left unclear their localization during later sporulation stages^23,28^. Here, our nanoscale analysis revealed an unexpected localization pattern at intermediate stages of TMR-star assembly (Fig. S6A). This prompted us to investigate in more detail the assembly dynamics of the early-synthesized inner coat proteins YutH and YisY, using single-sporangium time-lapse imaging. YutH and YisY were detected prior to the TMR-star signal, as expected for early-synthesized proteins (Fig. 3A and B and Fig. S8). However, rather than forming a continuous cap over the MCP forespore pole like the early-synthesized CotY (Fig. 1D), YutH and YisY initially localized as two distinct dots at the junction between the forespore sidewall and the MCP pole (Fig. 3A, time 0 and Fig. S8). The ‘two-dot’ appearance of YutH-GFP in 2D midplane sections was previously suggested to reflect a ring-like localization^28^. Sectioning of a ring at the midplane makes it appear as two dots spaced of the ring diameter. The YutH- and YisY-GFP signals then started spreading toward the midspore region but, instead of progressively encasing the entire forespore surface, they transiently covered the midspore region, before splitting in a ‘four-dot’ localization pattern. Notably, they disappeared from the midspore region when the TMR-star appeared and began to spread bidirectionally across this region (Fig. 3A and B and Fig S8). Strikingly, the GFP signals of both YutH and YisY were observed at the leading edge of the TMR-star front of extension appearing as though they were being ‘pushed’ forward by it (Fig. 3A and B). The distinctive localization of YutH and YisY on either side of the TMR-star signal was confirmed in snapshots of cells collected 5.5 to 7 hours after resuspension in sporulation medium (Fig. S6A). The same pattern was also observed for YsxE, the paralogue of YutH (Fig. S6A). In agreement with these observations, Costes PCC between YutH-GFP and TMR-star signals measured at the single-sporangium level over 200 minutes from TMR-star signal detection showed very transient and slight colocalization, only when TMR-star signal was first detected (t0 in Fig 3C) and again when TMR-star completely covered the forespore surface (t200 min in Fig 3C). Kymograph analysis of GFP/TMR-star signals over the entire forespore periphery further confirmed the intricate deployment of YutH-GFP and TMR-star signals, with YutH migrating bidirectionally at the leading front of TMR-star at a rate of 3.9 ± 1.2 nm/min (n = 32) (Fig. 3D). In contrast, CotY-GFP deployment was not coordinated with TMR-star labelling and proceeded mainly unidirectionally from the MCP towards the opposite MCD forespore pole at a rate of 4.5 ± 1 nm/min (n=11) (Fig. 3D).

### Displacement of YutH and deposition of TMR-star are spatiotemporally coordinated

Extensive displacement of molecules on the forespore surface has not been previously described in spore-forming bacteria. The dynamic redistribution of YutH and YisY along with their apparent anticorrelation with the TMR-star signal over time, prompted us to investigate their choreography in more detail. To this end, we collected cells expressing the YutH-GFP fusion at various stages of sporulation, labeled them with TMR-star, and imaged them using brightfield and dual-color, three-dimensional (3D) lattice-SIM². YutH-GFP 3D-localization patterns were classified relative to TMR-star localization to generate a virtual 4D-sequence of their dynamics (Fig. 4A). In top views of maximum (x,y) projections, YutH-GFP initially localized as a thin band at the junction between the midspore region and the MCP pole of the forespore, before TMR-star was detected (Fig. 4A, ‘x,y’, panel i). In front views of the corresponding maximum (y,z) projections, YutH-GFP appeared as an irregular ring (Fig. 4A, ‘y,z’ panel i). By imaging vertically immobilized cells, we further confirmed that YutH-GFP assembled as a ring with some density variation along its circumference (Fig. 4B, panel i). The timing and positioning of the YutH ring are similar to those recently described for the ring formed by the MucB/RseB-like protein SsdC^51^ (formerly YdcC) suggesting that YutH may associate with other proteins in this forespore region. Consistently, we observed similar polar ring localization for the YutH paralog YsxE and for YisY, which contrasts with the “typical” cap localization formed by CotY-GFP. When forespore refractility developed, but before TMR-star signal was clearly detected, the YutH ring expanded laterally (Fig. 4A, ‘Expansion’, ‘x,y’ panel ii) and appeared disrupted in different locations along its circumference (Fig. 4A, ‘y,z’ and Fig. 4B, panel ii, white arrowheads). Strikingly, when the TMR-star signal appeared in the mid-spore region and expanded bidirectionally towards the poles, YutH-GFP was progressively excluded from the zone labeled by TMR-star and displaced towards the two poles at the leading front of the TMR-star signal (Fig. 4A, ‘x,y’ panels iii-v, ‘Splitting’ and ‘Migration’). Accordingly, TMR-star was observed in regions devoid of YutH-GFP in cross-sections of vertically immobilized cells (Fig. 4B, panels iii-vi). Ultimately, TMR-star covered the entire forespore surface and YutH-GFP was restricted to the polar tips (Fig. 4A, panels vii, ‘Closure’). In conclusion, our reconstituted 4D-sequence revealed dynamic redistribution of YutH-GFP along the forespore cylinder during late stages of sporulation, spatiotemporally intertwined with TMR-star deposition.

**Figure 4.**
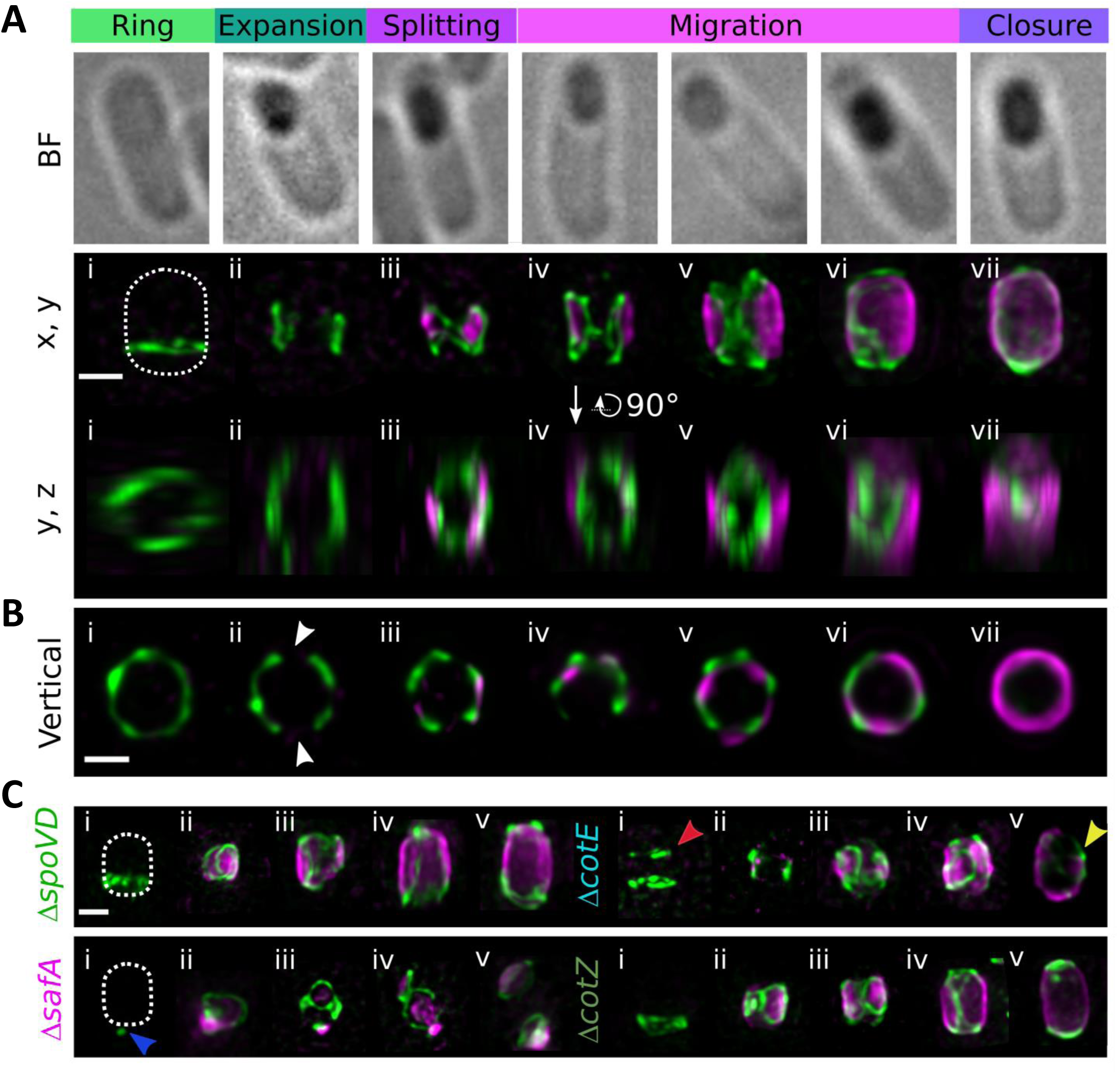
Redistribution of YutH is spatiotemporally linked to TMR-star deposition. **(A)** Brightfield (BF) images of representative sporangia and corresponding 3D lattice-SIM² reconstructions. Maximum (x, y) and (y, z) projections showing various super-resolved localization patterns of YutH-GFP (PE479, green) and TMR-star (magenta) observed during a sporulation of *B. subtilis.* The distinct stages of YutH-GFP virtual localization sequence are indicated on the top. The position of the FS is indicated by a white dotted line. **(B)** Representative 2D lattice-SIM² micrographs of vertically oriented sporangia, classified according to the progression of the TMR-star signal. White arrowheads point to gaps observed in the YutH ring just before TMR-star signal detection. **(C)** Representative 3D lattice-SIM² maximum (x, y) projection of YutH-GFP and TMR-star in the indicated genetic mutant background (Δ*spoVD*, RCL1697; Δ*safA*, RCL1698; Δ*cotE*, RCL1699; Δ*cotZ*, RCL1700). Blue arrowhead points to the abnormal localization of YutH in Δ*safA* non-refractile sporangia. Red and Yellow arrowheads point to the abnormalities in YutH localization observed in the Δ*cotE* mutant background. The position of the FS is indicated by a white dotted line in left panels. **(A-C)** Sporangia from at least two independent experiments, were collected between 5.5 and 7 hours after resuspension in sporulation medium. The various YutH-GFP localization patterns were ordered based on the extent of forespore coverage by the TMR-star signal, generating a virtual 4D super-resolved assembly dynamics sequence. Scale bars: **(A)**, 1µm; **(B and C)**, 500nm.

### Outer coat development influences YutH assembly dynamics

We next analyzed the 3D-localization of YutH in mutants lacking the morphogenetic factors SpoVD, SafA, CotE or CotZ, which involved in formation of the cortex, the inner coat, the outer coat and the crust, respectively. While it is known that YutH abnormally accumulates towards the MCP pole in absence of SafA^22^, we sought to determine whether the distribution dynamics of YutH is affected at later stage in absence of SafA and whether underlying (cortex) or overlying (outer coat and crust) coat layers might also affect YutH localization and/or dynamics. As previously described^22^, in the absence of SafA, YutH-GFP failed to initially assemble into its characteristic ring structure and instead accumulated near the MCP pole of the forespore (Fig. 4C, *ΔsafA* panel i, blue arrowhead). Interestingly, the TMR-star signal was also initially detected near the MCP pole, overlapping with YutH-GFP (Fig. 4C, *ΔsafA* panel ii), suggesting that the factor(s) required for YutH localization similarly influence the localization of the TMR-star substrate within the inner coat. Some aberrant migration of the YutH-GFP signal correlated with partial expansion of the TMR-star signal was observed in this region but this did not extend across the entire forespore surface (Fig. 4C, *ΔsafA* panels iii-v). No defects in YutH-GFP localization or dynamics were observed in the *ΔspoVD* and *ΔcotZ* mutants (Fig. 4C). In contrast, in the *ΔcotE* mutant, a second smaller YutH-GFP ring formed near the MCD pole of the forespore before becoming refractile (Fig. 4C, *ΔcotE* red arrowhead in panel i), suggesting that CotE, or a CotE-dependent (and CotZ-independent) protein, prevents YutH-GFP localization at the MCD pole. When TMR-star was detected in the *ΔcotE* mutant, we observed abnormal splitting and migration of YutH-GFP (Fig. 4C, *ΔcotE* panels ii-iv). These defects did not prevent complete deposition of the TMR-star signal over the forespore surface but impaired the final localization of YutH-GFP at the forespore polar tips (Fig. 4C, *ΔcotE* yellow arrowhead in panel v). Taken together, these results provide further evidence for a functional link between YutH-GFP and TMR-star assembly dynamics and suggest that the formation of the outer coat impacts the assembly of inner coat proteins.

### YutH displacement depends on late-synthesized factors

Although the expression of *yutH* and its paralog *ysxE* depends on the mother cell early transcription factor σ^E^, and not on σ^K5,52^, a previous study suggested that localization of YutH and YsxE depend on the late transcription factors σ^K^ and GerE^23^. To investigate the possible contribution of σ^K^ and GerE to the localization dynamics of YutH, we monitored the YutH-GFP fusion in the *ΔsigK* and *ΔgerE* mutants. In the *ΔsigK* mutant, TMR-star was not detected as shown above (Fig. 2F and G) but the YutH- ring initially formed and expanded correctly (Fig 5A, *ΔsigK* panels i and ii), However, YutH-GFP eventually spread across the entire forespore surface (Fig. 5A, *ΔsigK* panel iii), unlike in the WT background, where the expanded YutH-GFP ring split into two rings that migrate toward the poles upon TMR deposition (Fig. 4A and 5A, WT panel ii). Spreading of YutH-GFP over the surface of *ΔsigK* mutant refractile forespores correlated with a strong increase in GFP signal intensity relative to WT spores (Fig. 5B). In the *ΔgerE* mutant, where TMR-star signal over the forespore surface is strongly reduced (Fig. 2F and G), the initial YutH- GFP ring also formed correctly (Fig 5A, *ΔgerE* panel i) and ring splitting was observed (Fig. 5A, *ΔgerE* panel ii). However, migration of YutH-GFP signal towards the poles appeared strongly impaired and most of YutH-GFP molecules appear blocked in the midspore region (Fig. 5A, *ΔgerE* panel iii). Importantly, in *ΔgerE* refractile sporangia, where YutH migration was almost completely aborted, the level of YutH-GFP fluorescence is similar to the WT condition (Fig. 5B). This latter result supports a model in which a σ^K^ and GerE-dependent factor(s) drive(s) the extensive redistribution of YutH molecules over the forespore surface.

**Figure 5.**
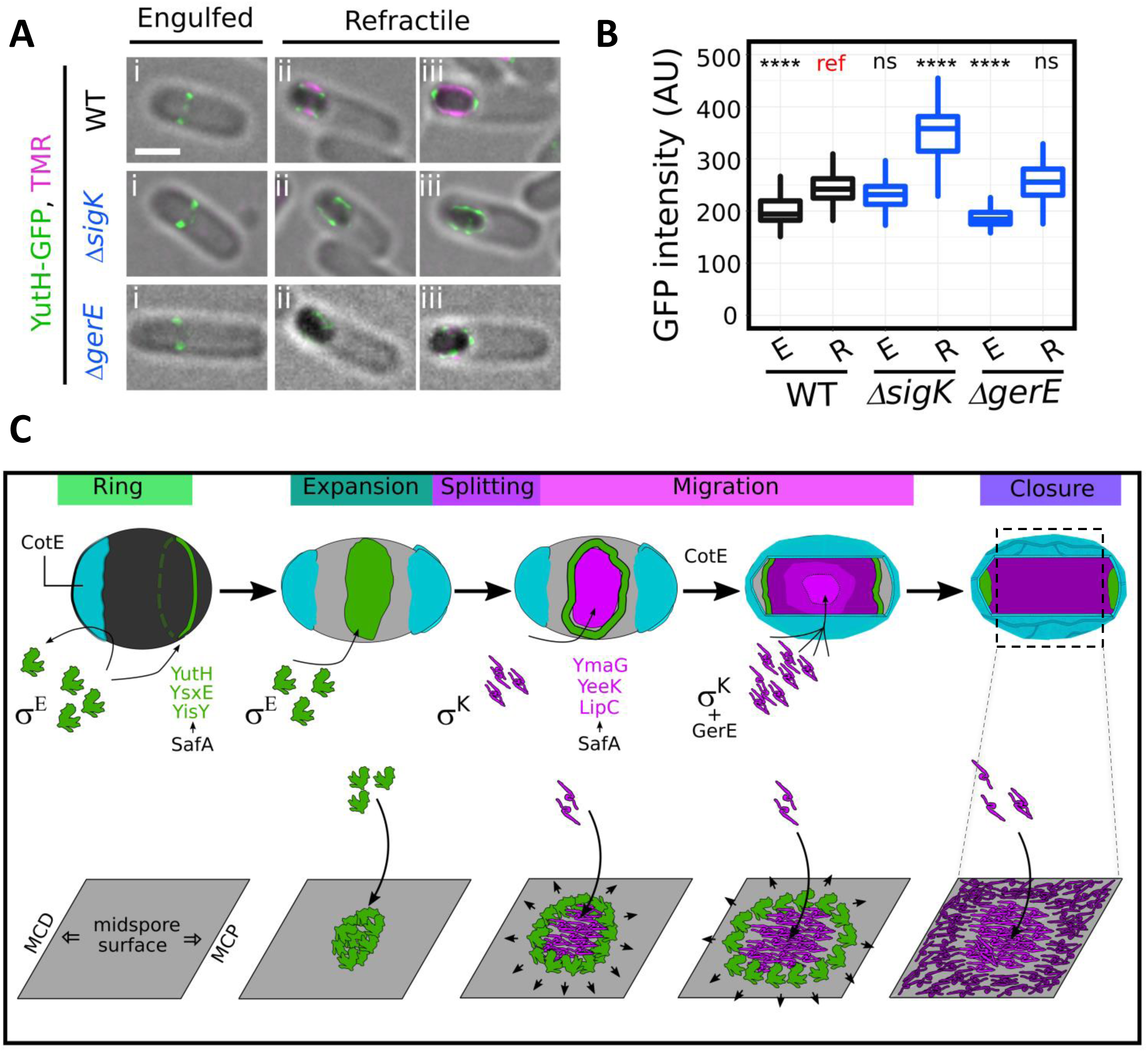
**Assembly of late-synthesized inner coat proteins in the midspore region leads to the relocation of pre-assembled material toward the forespore poles**. **(A)** Representative sporangia expressing YutH-GFP (in green) in the indicated genetic background (WT, PE479; Δ*sigK*, RCL1720; Δ*gerE*, RCL1696) collected between 5.5 and 7 hours after resuspension and having just completed engulfment (“Engulfed”) or showing a refractile forespore (“Refractile”). For the WT condition, a representative non-refractile sporangium showing the characteristic YutH ring at the junction between the MCP pole and the midspore region (i), refractile sporangium showing YutH migration (ii) or final YutH localization as two polar tips (iii) are shown. For mutant sporangia, the prevalent localization patterns are shown. Scale bar, 1 µm. **(B)** Quantification of YutH-GFP fluorescent signal intensity measured in corresponding pseudo-widefield images over the entire forespore surface in the indicated conditions (n>38 per condition, E: “Engulfed”; R: “Refractile”; A.U: Arbitrary Units). The non-parametric Mann–Whitney U test was used with the signal in WT refractile sporangia as a reference (ref); ns: not significant; ****P˂0.00005. **(C)** Model of inner coat (IC) development and YutH redistribution on the FS surface. Driven by σ^E^ synthesis and directed by SafA, YutH and associated early-synthesized IC proteins (YsxE and YisY), initially assemble as a ring-like structure at the junction between the FS MCP pole and the midspore region (“Ring” stage, see panel i in **A**). Assembly of CotE-dependent (and CotZ-independent) molecules towards the MCD pole (cyan) prevents the formation of a second ring. Note that CotE-dependent material is also present at the MCP pole but is not depicted here. Before σ^K^ activation, newly synthesized YutH molecules spread across the FS surface, expanding the ring-like structure towards the midspore region (“Expansion” stage). Following σ^K^ activation and dependent on SafA, late-synthesized IC proteins (YmaG, YeeK and LipC, in magenta) are produced and incorporated at the midspore region (“Splitting” and “Migration” stages), likely favored by the thinner outercoat in this area. As these late-synthesized IC proteins spread bidirectionally over the spore surface, they displace YutH towards the FS poles (“Migration” stage). The newest synthesized material (light pink), supported by GerE, integrates at the midspore region, pushing pre-assembled material (dark pink) towards the FS poles. Upon arrest of coat proteins production/integration, YutH is restricted to the FS pole tips (“Closure” stage). Migration of YutH molecules is linked to a CotE-dependent (and CotZ-independent) factor(s). Bottom panels show 2D projections of the midspore surface, illustrating the progressive waves of IC proteins deposition and the subsequent redistribution of pre-assembled material.

## Discussion

Here, by taking advantage of sporulation occurring in microcolonies, which tolerate repeated and intense laser exposure, we successfully combined prolonged single-cell time-lapse imaging with SIM to study coat formation in *B. subtilis*. Notably, by combining lattice-SIM illumination with the dual iterative deconvolution SIM² algorithm, we achieved a lateral resolution of approximately ∼70 nm, enabling us to distinguish the different coat sublayers. Furthermore, we demonstrate that TMR-star is a reliable maker for assessing the state of inner coat development in *B. subtilis*. The stable and linear binding of TMR-star to the developing inner coat allows to classify the localization patterns of virtually any protein in sporangia with a refractile forespore, and enables precise kinetic analysis of cells collected and imaged at late stages of sporulation. This approach appears particularly useful for the study of proteins with a nonlinear deployment, such as the early-synthesized inner coat proteins YutH, YsxE and YisY, and for generating a faithful 4D-sequence of coat assembly. Our time-lapse super-resolved approach could be used to explore new aspects of coat development that remain inaccessible when analyzing cell populations collected at different times after resuspension. For example, we showed that crust assembly (as judged by CotY-GFP incorporation) is completed later than completion of inner coat assembly (judged by TMR-star incorporation). Paradoxically, CotY-GFP deployment starts earlier than TMR-star binding (and subsequent initiation of YutH-GFP migration). This apparent paradox can be resolved by the bidirectional displacement of YutH/TMR-star, whereas CotY assembly proceeds mainly unidirectionally from the MCP pole. Using TMR-star labelling combined with single-cell Lattice-SIM² time-lapse imaging, we recently discovered that KapD, a ribonuclease localized in the spore crust, is also redistributed on the forespore surface upon activation of σ^K^ (D’halluin et al., accepted). Collectively, these results reveal that the dynamic relocation of pre-assembled early-synthesized coat proteins on the forespore surface is not restricted to the inner coat formation but can also occur in other layers of the *B. subtilis* coat, and possibly in other (if not all) spore-forming bacteria.

We also showed that the inner coat layer assembles with distinct dynamics relative to the external coat layers (the outer coat and crust). Specifically, late-synthesized inner coat proteins such as YmaG, LipC, and YeeK, initially localize to the midspore region. Interestingly, YppG, a late-synthesized protein located in the undercoat, underlying the inner coat, also assembles first in the midspore region (Fig. S6A). Thus, initial assembly in the midspore region appears as a common feature shared by proteins from innermost coat layers (inner coat and undercoat). Importantly, this behavior is reminiscent of coat material deposition previously reported in *B. cereus* species^35,37–39^. Furthermore, we showed that TMR-star specifically labels the *B. subtilis* inner coat and that its localization depends on σ^K^, GerE and SafA, as reported in *B. cereus* species^39,53^. This suggests that the inner coat of *B. subtilis* and the coat layer of *B. cereus* species share structural similarities and that the use of TMR-star can most likely be extended to studies of coat development in other less studied and less genetically tractable *Bacillaceae* species.

Another major observation is the dynamic relocalization of the early-synthesized inner coat proteins YutH (and to lesser extent YisY) which is spatiotemporally synchronized with TMR-star deposition, generating unique localization patterns over the forespore surface. This particular dynamic is likely shared by YsxE, the YutH paralog, which exhibits similar localization patterns (Fig. S6A) and similar genetic dependencies for its assembly^23^. Interestingly, YutH orthologs are among the best conserved coat proteins among *Bacillaceae*^4,29,54^ suggesting that this mechanism is largely conserved. We propose a model in which YutH, YsxE and YisY initially form a ring structure, likely with other early-synthesized sporulation proteins such as SsdC^51^, at or near the junction between the MCP pole and the spore sidewall (Fig. 5C). Then, they spread towards the midspore region, likely driven by protein neo-synthesis occurring after engulfment completion, before splitting into two rings that migrate towards the forespore pole tips (Fig. 5C).

The main question arising from these observations is: what mechanism drives this unusual molecular displacement on the forespore surface? Our findings provide evidence that YutH and YsxE are displaced towards the forespore poles as late-synthesized coat material assembles in the midspore region (Fig. 5C). The main result supporting this model is that synthesis of YutH and YsxE does not depend on late transcription factors but their relocalization is strongly affected in the absence of σ^K^ or GerE. In absence of σ^K^ we found an apparent increase in YutH incorporation on the forespore surface, indicated by the increased GFP intensity measured in Δ*sigK* refractile sporangia. This result mirrors the increased expression of the YutH paralog YsxE previously reported in the *ΔsigK* mutant^52^, which was attributed to the absence of σ^E^ shutdown in the mother cell, leading to abnormal production of early-synthesized proteins at late stages of sporulation^55,56^. In absence of GerE, the overall levels of YutH were similar to the wild-type. Since *sigK* expression is repressed by GerE^57^, this result further supports that YutH is only produced by σ^E^. Importantly, in the absence of GerE, the migration of YutH was severely impaired indicating that a GerE-dependent factor is responsible of YutH molecules relocation from the midspore region to the poles. To test this model, we also analyzed YutH-GFP and TMR-star localization in several mutants deficient for late-synthesized inner coat proteins. However, we failed to detect any defects indicating either redundancy between these factors or a yet uncharacterized, σ^K^, GerE, and SafA-dependent factor, likely conserved with *B. cereus* species, that allows TMR-star binding and is responsible for YutH displacement.

Another intriguing question braising from our observations is: why are some late-synthesized coat proteins targeted to the midspore region rather than to the forespore poles? Targeting of late-synthesized outercoat proteins (CotQ, CotS, YtxO) to the forespore pole(s) was clear in previously reported diffraction-limited images^23^. Here, we confirmed a similar behavior for SpsI, a late-synthesized outer coat protein^58^ and for CgeA, a late-synthesized crust protein^17,59^ (Fig. S6A). In contrast, the late-synthesized inner coat proteins YmaG, LipC and, to a lesser extent, YeeK were initially detected along the long axis of the forespore similar to TMR-star labelling. This observation suggests that the long axis of the forespore is the primary targeting site for several proteins.. In *B. cereus* species, deposition of coat material towards the midspore region is linked to the presence of the exosporium cap at the forespore pole, interfering with deposition of coat material at this position by an unknown mechanism^35,37,39^. Based on the similarities we highlighted between the development of the inner coat of *B. subtilis* and the coat of *B. cereus* species, we propose that, similarly, late-synthesized inner coat proteins cannot access the forespore poles in *B. subtilis* because of the accumulation of outer coat/crust material at the poles (Fig. 5C). Supporting this idea, in vertical imaging experiments, the outer coat layer (visualized via SpsI-GFP localization) was juxtaposed to the inner coat layer (as judged by TMR-star localization) towards the midspore region but appeared external to it at the poles. This finding echoes prior TEM studies that noticed differences in outer coat thickness between the polar regions and the midspore region^40^. We did not observe any effect of a *cotZ* deletion (essential for crust formation, the equivalent of the exosporium) on TMR-star localization or YutH assembly dynamics. Strikingly, however, in a Δ*cotE* strain, YutH assembly dynamics were modified with, i) a second ring of YutH seen towards the MCD pole at an early stage of sporulation and ii) an abnormal YutH migration viewed at late stages of sporulation. Together these results suggest that across *Bacillaceae*, the outermost spore layers preclude access of the forespore poles to late-synthesized proteins destined to the innermost layers, which are consequently incorporated at the midspore region (Fig. 5C). Finally, since i) forespore poles are the last regions to be covered by late-synthesized inner coat proteins, and ii) the YutH molecules, which synthesis appears not supported by GerE, are progressively displaced from the midspore towards the forespore poles, we propose that late-synthesized proteins from the innermost coat layers are continuously incorporated towards the midspore region (light pink in Fig. 5C), pushing preassembled material (dark pink in Fig. 5C).

In conclusion, our study provides novel approaches for in-depth study of dynamic processes occurring during *B. subtilis* sporulation, with a particular focus on the latest stages of spore development. These stages remain largely unexplored due to the limitations of light microscopy and absence of a suitable physiological dye to accurately label these stages. Furthermore, our study bridges the gap between models of proteinaceous layers formation among distant *Bacillaceae* species; *B. subtilis* and *B. cereus* species, and highlights new mechanisms involved in the formation of the multilayered *B. subtilis* coat.

## Methods

### Bacterial strains and culture conditions

All *B. subtilis* strains analyzed in this study are congenic derivatives of the standard parental strain PY79^60^ and are listed in Table S1. Fluorescent fusion to the Rny single-pass membrane was synthesized and sequenced by Genecust, cloned using BamHI/EcorI restriction sites in pDG1730^61^ and integrated at *amyE* locus under control of a synthetic promoter (see sequence in Fig. S4) to generate strain RCL1736. Gene deletion strains were either previously constructed or obtained from the whole genome deletion library (BKE or BKK strains^62^), the genomic DNA was extracted using the High Pure PCR Template Preparation Kit (Roche), transferred in PE479 strain by natural transformation and selected according to the antibiotic resistances. Strains were struck out onto LB plates from frozen glycerol stocks and incubated at 30°C overnight. Single colonies were then used to inoculate a 3mL volume of 1/4 diluted LB overnight at 30°C. Overnight cultures were diluted 1/20 in 1/4 diluted prewarmed LB and after growing cells at 37°C to OD600 (optical density at 600 nm) ∼0.3-0.5, sporulation was induced by resuspension in A+B medium according to the method described by Sterlini and Mandelstam^63^ at 37°C.

### Lattice-SIM imaging

Samples (800 µL) were withdrawn from cultures in A+B at selected times during sporulation or after 5.5 hours for time-lapse imaging. Cells were collected by centrifugation (9,000×g, 2 min), resuspended in 100 µL of supernatant and labeled by incubation with SNAP-cell TMR-Star (New England Biolabs) at a final concentration of 250 nM for 30 min at 37 °C in the dark. For time-lapse imaging, the TMR-star-stained cells suspension was directly applied onto a gene frame containing a 1.6% agarose pad in A+B media (1 to 4 different strains were visualized simultaneously using small sections of the pad). For snapshots and 3D acquisition of cells collected during the time-course of sporulation, the TMR-star labeling step was followed by 2 washes with 1 mL of PBS. All experiments were done at 30°C in a temperature-controlled chamber. For lattice-SIM² time-lapse imaging of sporulation in microcolonies, we adapted our protocol from a previously published one^64^. The mounted slides were incubated overnight in the dark at 30°C allowing to trigger microcolonies formation on the agarose pad. We observed a reduced proportion of pre-germination events when sporulation occurs in these microcolonies compared to cells sporulating in liquid medium that are directly imaged after TMR-star labelling by lattice-SIM² (Fig. S1A). Samples were imaged with an Elyra 7 AxioObserver (Zeiss) microscope equipped with an alpha Plan-Apochromat 63x/1.46 Oil Korr M27 objective and a Pco. edge 5.5 camera, using 488 nm (100 mW) or 561 nm (100 mW) laser lines. For each lattice-SIM acquisition, the corresponding grating was shifted 15 times for snapshots or 13 times for 3D and time-lapse acquisitions to reduce light exposure. The grid periods used were 28 mm or 34 mm for acquisitions with the 488 nm or 561 nm lasers respectively. For 3D-acquisitions, 29-31 sections of 0.1 µm were acquired with “center” function of Zen software. For detection of the GFP signals, the laser was settled at 30% of maximal power with 30 ms exposure per frame for CotY-, YutH-, YhaX-, SpsI-, CgeA-, YsxE-and Yeek-GFP fusions; 20% laser power and 20 ms exposure per frame for YisY-, YmaG-, YxeE-GFP and Nrny-sfGFP fusions; 30% laser power and 60 ms per frame for LipC-GFP; 40% laser power and 40 ms exposure for YppG-GFP and 10% laser power and 20 ms per frame for CotD-GFP. For snapshots and 3D imaging of cells collected during the time-course of sporulation, TMR-star signal was illuminated at 10% of maximal laser power using 20ms exposure per frame. For time-lapse analysis of YutH-GFP in various mutant context, 10% laser power and 50 ms exposure time were used. Brightfield snapshots of the area imaged were acquired before and after time-lapse imaging to confirm the success of sporulation (Example in Fig. S1A). Final lattice-SIM images were reconstructed using the ZEN software (black edition, 2012) and the dual iterative SIM² algorithm with settings previously described^65^, yielding a final pixel size of 16 nm for reconstructed images.

### Vertical cell imaging

For vertical imaging, silicon molds described elsewhere^65^ were prepared to print a field of microholes on agarose pads. These pads were prepared by pouring melted 6% agarose in A+B medium on slides, immediately covered with the silica molds to generate microholes. 6 μL of sporulating cells collected after hour 5.5 to 7 were deposited on agarose pads and left to rest 1-2 min before being centrifuged in a mini-centrifuge (10 s, 2000 g) and covered with 5µL of melted 1% agarose in A+B medium and a cover glass before imaging with lattice-SIM².

### Image processing and analysis

All image analyses were performed and micrographs were processed using the Fiji software (ImageJ, NIH). Brightness and contrast of representative cell images were adapted in Fiji and figures were compiled using Inkscape 0.92.5. (https://inkscape.org). At least three different microscopic fields were analyzed for each condition. Quality check is an important step in SIM images interpretation due to their susceptibility to artefact generation^66^. While SIMcheck, a commonly used plugin developed for detecting such defects, is highly effective^66^, it is unfortunately not yet compatible with lattice-SIM illumination patterns. To assess the quality of our lattice-SIM reconstructed images, we therefore used SQUIRREL, a tool designed to detect reconstruction artifacts in super-resolution images^67^. SQUIRREL computed two key metrics, the resolution scaled Pearson’s correlation (RSP) and the resolution scaled error (RSE), that report how well the reconstructed image correlates with the pseudo-widefield image (Fig. S9). For RSP, values closer to 1 reflect better the agreement. For RSE, the smaller the number, the better the agreement, 0 being a perfect agreement. Our analysis using SQUIRREL revealed a high RSP value (>0,9) and a low RSE value across the field of views analyzed, with low error levels near the sporulating cells (see inset in Fig. S9), confirming the absence of artefacts in reconstructed images. Sporulation in microcolonies imposes certain challenges for images post-processing and analysis. In addition to drift occurring during long time-lapse acquisition, cells dividing during microcolonies development eventually push and displace sporulating cells, rendering complex the alignment and correction steps. To circumvents these issues, we first performed a manual alignment step using a fixed object; an isolated spore. In contrast to fluorescent beads, isolated and immobilized spores showed a fluorescence of similar intensity to the one of studied objects (here the sporangia) optimizing SIM² reconstruction. Then, cells were aligned automatically using the “Image Stabilizer” plugin of Fiji with the TMR-star channel used to extract log of transformation that was applied to the green channel using “Log Image Stabilizer” plugin. Alignment-corrected images were then used to create representative montages, kymographs, 3D-maximym Z-projections and to perform colocalization analysis shown in this study. For colocalization analysis, when possible, multicolour SIM images were aligned with the “channel alignment” tool in Zeiss ZEN black software and parameters set to “Fit” and “Lateral”. In order to measure the Costes Pearson Correlation Coefficient (PCC) between the GFP and TMR-star signals, aligned images of each fluorescence channel for individual sporangia, from either time-course snapshots or time-lapse experiments, were analyzed with the Fiji plugin “JaCoP” v2.1.121 using the score corresponding to the “Costes automatic threshold”^49,50^. When no colocalization was found by the function, a score of 0 was attributed by the function. Kymographs were generated in Fiji using the ‘Kymograph builder’ plugin. Quantification of the GFP intensity in forespore of various genetic background was performed on pseudo-widefield images. A polygonal region of interest (ROI) was adjusted using the Fiji polygon tool on the periphery of the forespore surface and the mean intensity was recorded for the indicated numbers of sporulating cells.

### Statistics and reproducibility

Lattice-SIM data collection was performed at least twice independently in time-course experiment before being imaged once with Time-lapse. Statistical analyses were performed using Rstudio version 4.1.1 for PC. Data are represented with boxplots showing the interquartile range (25th and 75th percentile). The upper whisker extends from the hinge to the largest value no further than 1.5×IQR from the hinge and the lower whisker extends from the hinge to the smallest value at most 1.5×IQR of the hinge.

## Data Availability

The data that support this study are available from the corresponding author upon request.

## Aknowledgements

We thank the members of the ProCeD laboratory for their support and helpful discussions; Patrick Eichenberger and Adriano O. Henriques for providing strains; Nathalie Laforge, Philéas Larcher, Ingrid Adriaans and Aurélien Barbotin for help with early works on this project; Cyrille Billaudeau for expertise in SIM acquisitions and analysis of reconstructed data and Adriano O. Henriques for critical reading of the manuscript. This project has received funding from the Agence Nationale de la Recherche (ANR-21-CE12-0018 to R.C.-L. and C.C.) and from the European Research Council (ERC) under the Horizon 2020 research and innovation program (ERC-2017-CoG-772178 to R.C.-L.).

## Authors contributions

A.L. and R.C.L. designed research. A.L. and D.J performed experiments. A.L., D.J. and R.C.L. analyzed data. A.L. wrote the manuscript, with input from all authors; A.L. and R.C.L. revised the manuscript. C.C and R.C.L acquired funding.

---

**Table S1.**
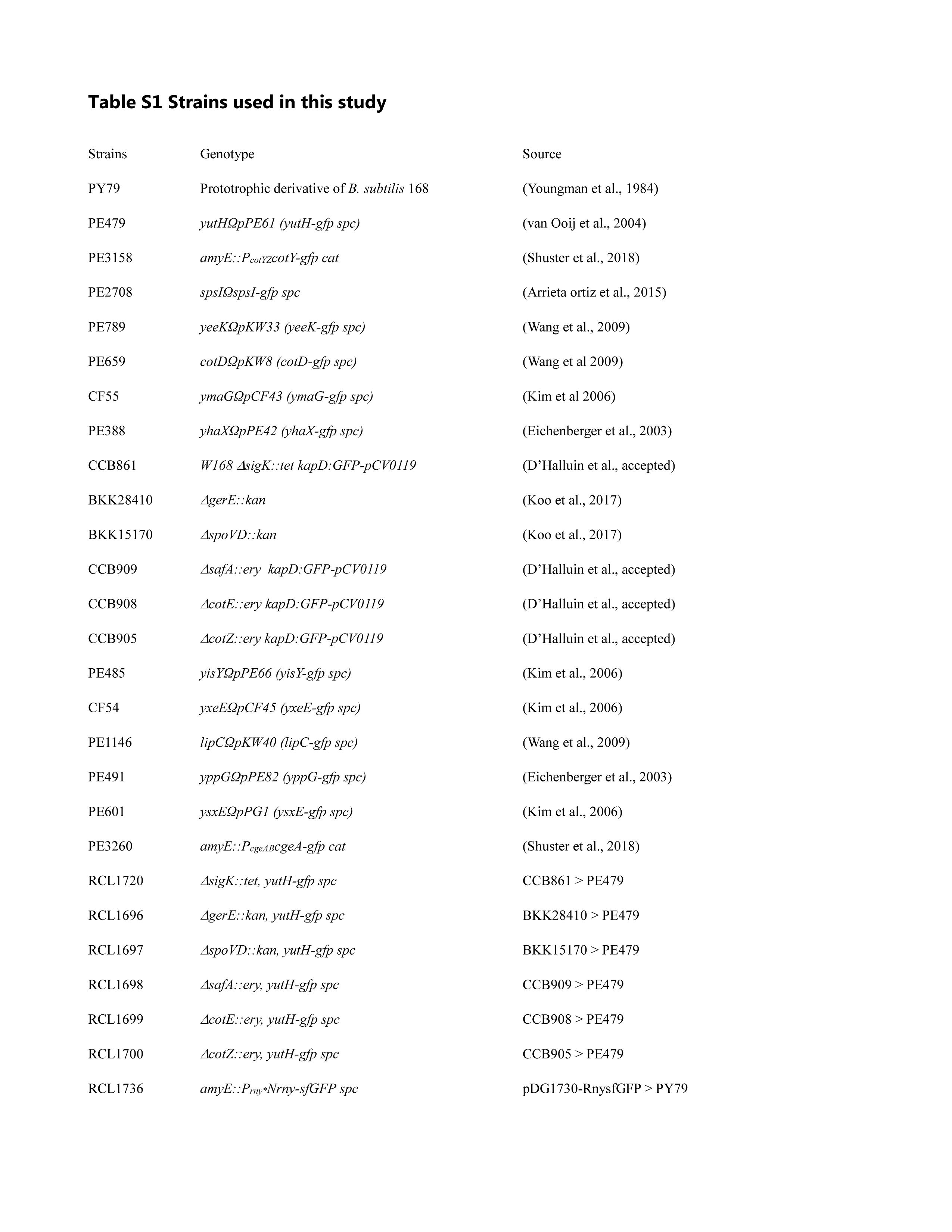
Strains used in this study.

## Supplemental Figure Legends

**Figure S1. Workflow of lattice-SIM² imaging of *Bacillus subtilis* sporulation. (A)** Schematic of the experimental procedures used for time-course (top panels) or time-lapse (bottom panels) analysis of *B. subtilis* sporulation. For time course experiments, cells resuspended in sporulation medium (SM) are harvested at different time points during the time-course of sporulation. Previously, a virtual sequence of coat proteins assembly was derived from fluorescence images of sporangia population. Here, we labeled the cells with TMR-star, and imaged immediately with 3D lattice-SIM². For time-lapse at the single-sporangium level, we performed lattice-SIM microscopy of cells sporulating in microcolonies. Cells were collected 5.5 hours after resuspension, labelled with TMR-star, mounted on an agarose-coated microscope slide and incubated overnight at 30°C. Non-sporulating cells multiply, forming microcolonies on the agarose pad. Sporulation occurring in those microcolonies is highly efficient, as indicated by development of refractile forespores, despite repeated and intense laser exposure. Up to four strains can be imaged in parallel for up to 12 hours. Following SIM² reconstruction and post-processing corrections of Lattice SIM acquisitions (see Methods), quantitative parameters such as protein encasement speed can be extracted. Furthermore, by monitoring the single-sporangium 2D sequences of GFP-tagged coat protein assembly (488 nm laser) relative to the TMR-star signal (561 nm laser), a super-resolved 4D assembly sequence can be proposed. **(B)** A negligible fraction of sporangia exhibited premature germination event in cells sporulating in microcolonies compared to cells sporulating in liquid medium and collected for immediate visualization. The graph shows the percentage of premature germination events (“Preger.”) among the population of refractile sporangia (“Refrin.”). Experiments were performed at least 3 times for the two conditions.

**Fig. S2. Assembly dynamics of CotY-GFP and TMR-star in a dual color Lattice-SIM² single-sporangium movie.** Single-sporangium time-lapse showing the co-localization of CotY-GFP (green, PE3158) and TMR-star (magenta) in strain PE3158 by Lattice-SIM² super-resolved microscopy. Images are shown every 12 minutes over 10 hours. Green and Magenta boxes indicate the first frame of CotY-GFP and TMR-star detection, respectively. White areas (outlines) highlight the expansion of GFP and TMR-star signals on the forespore cross-section. Scale bar, 500nm.

**Figure S3. Effect of expression of GFP fusions to coat proteins on spore dimensions and TMR-star signal. (A)** Schematic indicating the timing of synthesis (early or late) and the localization of various GFP fusions to indicated coat proteins (wo, PY79; YhaX, PE388; YppG, PE491; LipC, PE1146; YmaG, CF55; CotD, PE659; YeeK, PE789; YutH, PE479; YsxE, PE601; YisY, PE485; SpsI, PE2708; CotY, PE3158; CgeA, PE3260), according to Driks and Eichenberger (2016). *, Note that LipC localization was unclear in this previous study (magenta dotted boxplots in B-D), being classified either as an under coat or as an inner coat protein (Driks and Eichenberger, 2016). Here, the extensive colocalization of LipC-GFP with TMR-star, similar to other late-synthesized inner coat proteins and in contrast to the under coat YppG fusion (see Fig. S6), supports that LipC is an inner coat protein. **(B)** Quantification of the sporangium length (from MCP pole to MCD pole) based on the TMR-star signal in sporangia expressing the indicated GFP-fusions. At least 22 sporangia per condition were analyzed, except for the YppG condition (n=12). A schematic of a sporangium with a dotted line indicating the region used for quantification is shown at the top. wo, without GFP-fusion expressed. **(C)** Full Width at Half Maximum (FWHM) quantification of TMR-star signal in refractile sporangia expressing the indicated GFP-fusions. At least 12 sporangia were analyzed per condition. The non-parametric Mann–Whitney U test was used with “wo” as a reference (ref); ns: not significant. **(D)** FWHM quantification of GFP-coat fusions signal in refractile sporangia expressing the indicated GFP-fusions. At least 12 sporangia were analyzed per condition. The non-parametric Mann–Whitney U test was used with strain expressing YutH-GFP as a reference. ns: not significant; *P<0.05; ****P<0.00005.

**Figure S4. Assessment of lattice-SIM² lateral resolution. (A)** DNA sequence of the Rny single pass N-terminal helix domain (amino acid 1-30) fused to sfGFP cloned into pDG1730 plasmid using BamHI/EcoRI sites and integrated at the *amyE* locus by a double recombination event (RCL1736). A synthetic promoter derived from the natural PymdA promoter was used (Guiziou *et al*., 2016). **(B)** Comparison of pseudo-widefield (pWF), lattice-SIM and lattice-SIM² image of strain RCL1736 expressing the Nrny-sfGFP fusion as a membrane reporter, collected 5.5 hours after resuspension in sporulation medium. Brighfield (BF) illumination allows to distinguish engulfing membranes. The magenta dotted line in the pWF panel indicates the region used for quantification in panel C. **(C)** Mean and standard deviation of the quantification of FWHM as a proxy for lateral resolution of Nrny-sfGFP membrane signal (n=15). The non-parametric Mann–Whitney U test was used with the “SIM” condition as a reference; ****, P<0.00005. Scale bar is 1µm.

**Figure S5. TMR-star colocalizes with the late-synthesized inner coat proteins YeeK and CotD. (A)** Representative micrographs showing co-localization of the late-synthesized inner coat proteins YeeK (strain PE789) and CotD-GFP (strain PE659) relative to TMR-star when it fully covers the forespore surface. Scale bar, 500 nm. Line profiles analysis (from pole to pole) show the superposition of GFP and TMR-star normalized signals. N.F.I, Normalized Fluorescence Intensity. **(B)** Distribution of Costes Pearson Correlation Coefficient (Costes PCC) between signals of TMR-star and GFP fused to the indicated coat proteins. The non-parametric Mann–Whitney U test was used with YmaG-GFP as a reference (ref); ns: not significant (at least 22 sporangia per condition).

**Figure S6. Analysis of various coat proteins localization at an intermediate stage of TMR-star assembly. (A, B)** Representative super-resolution micrographs showing localization of the indicated early-synthesized or late-synthesized coat protein fused to GFP (YhaX, PE388; YppG, PE491; LipC, PE1146; YmaG, CF55; CotD, PE659; YeeK, PE789; YxeE, CF54; YutH, PE479; YsxE, PE601; YisY, PE485; SpsI, PE2708; CotY, PE3158; CgeA, PE3260), when TMR-star is only detected on the midspore region and not detected towards the forespore poles. (A) and control of YmaG-GFP localization in absence of TMR-star labelling (B). Sporangia were collected between hours 5.5 and 7 after resuspension in sporulation medium. Scale bar, 500 nm; BF, brightfield. **(C)** Distribution of Costes Pearson Correlation Coefficient (Costes PCC) between signals of TMR-star (only viewed in midspore region) and GFP fused to the indicated coat proteins over the entire FS surface. For the crust proteins CotY and CgeA, the software was not able to compute a Costes PCC because of a complete absence of correlation between GFP and TMR-star signals in these conditions. PCC=1 would indicate perfect colocalization; PCC=0, no colocalization; PCC>0.5 is considered as a significant colocalization (ref line). The non-parametric Mann–Whitney U test was used with LipC-GFP as a reference (ref); ns, not significant; ****, P<0.00005 (at least 16 sporangia per condition).

**Figure S7. Single-sporangium dual color Lattice-SIM² time-lapse showing the localization of the late-synthesized inner coat proteins YmaG, LipC and YeeK relative to the TMR-star signal.** Original time intervals of the time-lapse acquisition are indicated. Green and magenta squares indicate the first frame of GFP-coat fusion (LipC, PE1146; YmaG, CF55; YeeK, PE789)and TMR-star detection, respectively. For YeeK-GFP, the area of increased intensity that followed the assembly of TMR-star is highlighted with a white outline. Scale bar, 500 nm.

**Figure S8. Single-sporangium dual color Lattice-SIM² time-lapse showing the localization of the early-synthesized inner coat proteins YutH and YisY relative to the TMR-star signal.** Original time intervals are indicated. Green and Magenta squares indicate the first frame of GFP-coat fusion (YutH, PE479; YisY, PE485) and TMR-star detection respectively. Scale bar, 500 nm.

**Figure S9. Control of Lattice-SIM² reconstructions using SQUIRREL analysis**. Sporulating cells expressing Nrny-sfGFP fusion (RCL1736) collected 5.5 hours after resuspension were imaged using lattice-SIM and reconstructed with the SIM² algorithm as described in the Methods section. A pseudo-widefield (pWF) image used as a ‘Raw’ image, a lattice-SIM² super-resolution reconstruction, a super-resolution image convolved with an automatically computed RSF (’Convolved’), and a quantitative map of errors between reference and convolved SR images (’Error map’; color bar indicates magnitude of the error) are depicted. The error maps show the resolution-scaled error (RSE), calculated between the blurred super-resolution lattice-SIM² image and the pWF image. The resolution-scaled Pearson coefficient (RSP) represents a normalized quality metric, showing high values and thus high-quality reconstructions for lattice-SIM² reconstructions. An inset showing two sporulating cells is shown in the bottom panels.

