## Supplemental Figures for "Real-Time Nanoscale Investigation of Spore Coat Assembly in *Bacillus subtilis*"

Figure S1

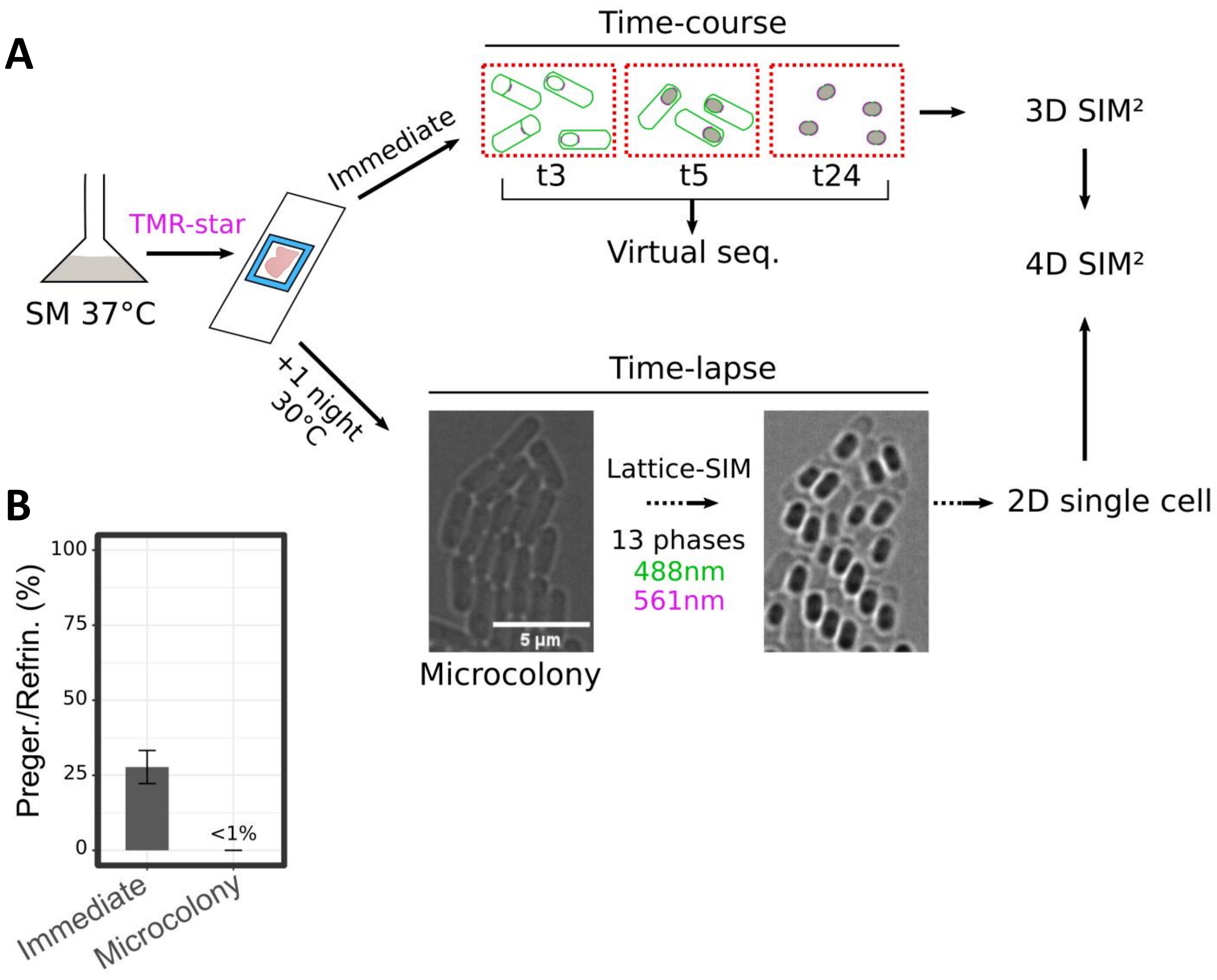
A
Time-course
Immediate
t3
t5
t24
3D SIM²
Virtual seq.
4D SIM²
TMR-star
SM 37°C
+1 night
30°C
Time-lapse
Lattice-SIM
13 phases
488nm
561nm
5 µm
Microcolony
2D single cell
B
Preger./Refrin. (%)
100
75
50
25
0
<1%
Immediate
Microcolony

Figure S2

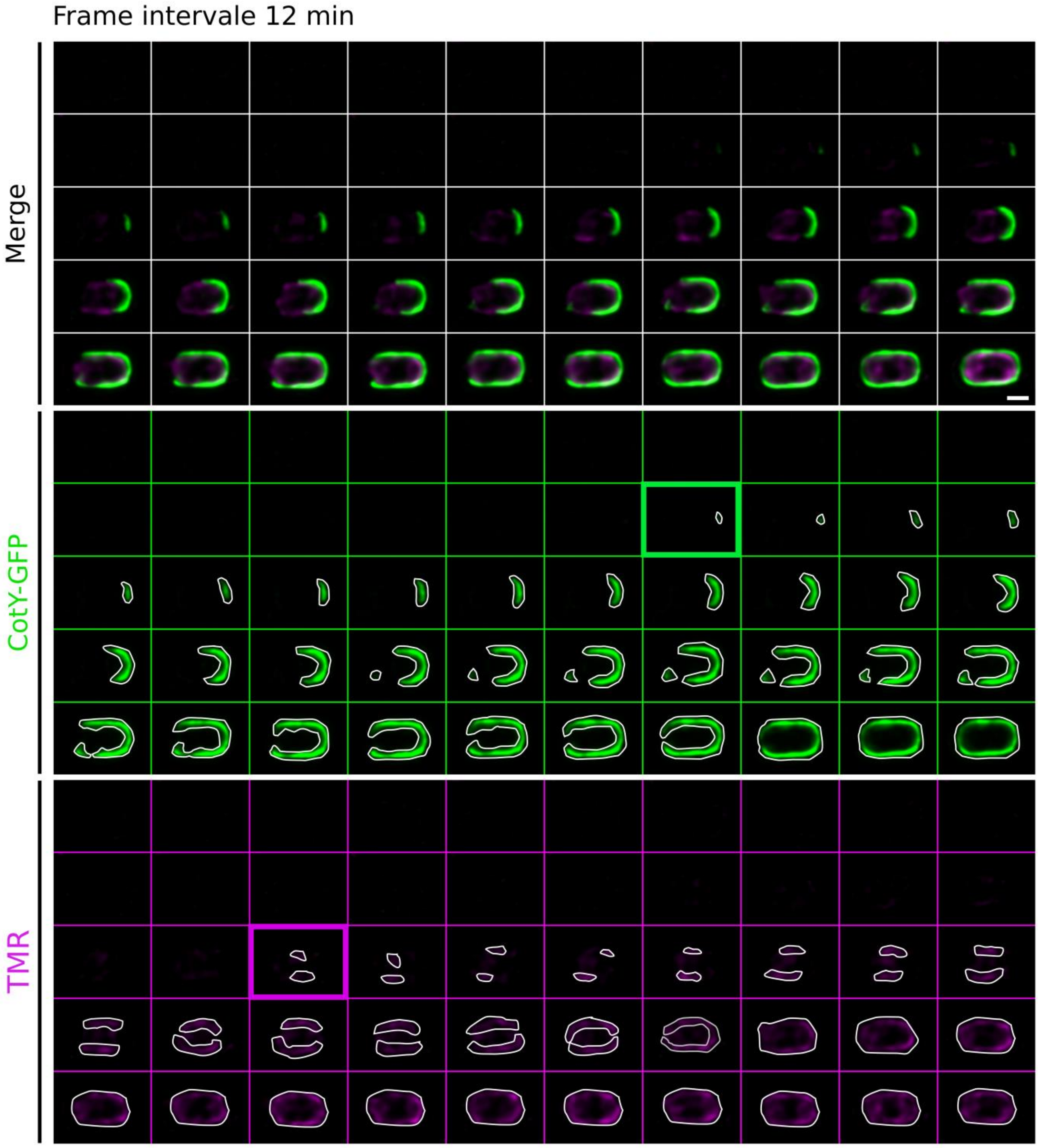

Figure S3

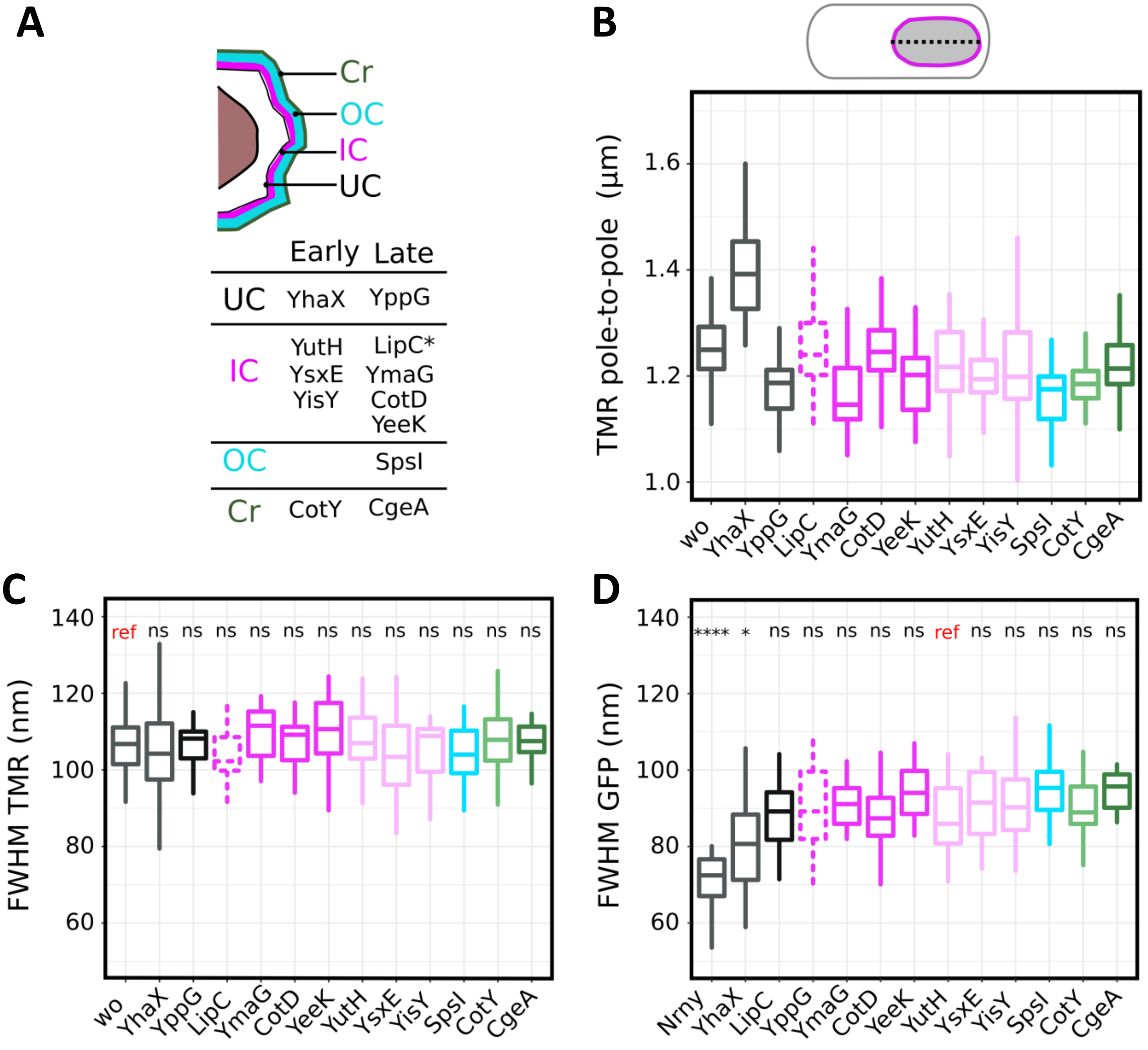
A
Cr
OC
IC
UC
Early Late
UC YhaX YppG
IC YutH YsxE YisY LipC* YmaG CotD YeeK
OC SpsI
Cr CotY CgeA
B
TMR pole-to-pole (µm)
1.0
1.2
1.4
1.6
wo YhaX YppG LipC YmaG CotD YeeK YutH YsxE YisY SpsI CotY CgeA
C
FWHM TMR (nm)
60
80
100
120
140
ref ns ns ns ns ns ns ns ns ns ns ns ns
wo YhaX YppG LipC YmaG CotD YeeK YutH YsxE YisY SpsI CotY CgeA
D
FWHM GFP (nm)
60
80
100
120
140
**** * ns ns ns ns ns ref ns ns ns ns ns
Nrny YhaX LipC YppG YmaG CotD YeeK YutH YsxE YisY SpsI CotY CgeA

**A**

BamHI-spacer-PymdA-**Nrny[1-30]**-sfGFP-spacer-EcoRI

```
GGATCCGATCCTGTATTACTATTCTTAGTTAAGATGGCAAGCTTGACAAGTATTTCCGACACATTTACA
ATGAAGTTGGAGAAAAGCTAGCGATTAACTAATAAGGAGGACAAACATGACCCCAATTATGATGGTTCT
CATCTCCATTTTGCTGATTCTACTCGGTTTAGTTGTTGGCTACTTTGTTCGTAAAACCATTGCCGAAAT
GTCAAAAGGAGAAGAACTTTTTACAGGTGTAGTACCTATCTTGGTTGAATTGGATGGTGATGTTAACGG
TCACAAATTTTCTGTACGTGGTGAAGGTGAAGGTGATGCAACTAACGGTAAATTGACACTTAAATTCAT
TTGTACAACTGGAAAACTTCCTGTTCCTTGGCCTACTCTTGTTACAACATTGACATATGGAGTACAATG
TTTTTCACGTTATCCTGATCATATGAAACGTCACGATTTTTTTAAATCTGCTATGCCAGAAGGTTATGT
ACAAGAACGTACAATTTCATTTAAAGATGACGGAACATATAAAACACGTGCTGAAGTAAAATTCGAAGG
TGACACTCTTGTTAATCGTATCGAATTGAAAGGAATCGATTTCAAAGAAGATGGTAACATTTTGGGACA
CAAACTTGAATACAACTTCAACTCTCATAATGTTTATATCACAGCTGACAAACAAAAAAACGGTATTAA
AGCTAATTTTAAAATTCGTCACAATGTTGAAGATGGATCTGTTCAATTGGCTGATCATTATCAACAAAA
TACACCAATCGGAGACGGACCAGTATTGCTTCCAGATAACCACTACCTTTCTACTCAATCAGTTCTTTC
AAAAGATCCTAACGAAAAACGTGACCATATGGTACTTCTTGAATTTGTTACAGCAGCAGGTATCACTCA
CGGTATGGACGAACTTTATAAATAAACTTTATCTGAGAATAGTCAATCTTCGGAAATCCCAGGTGGAAT
TC
```

**B**

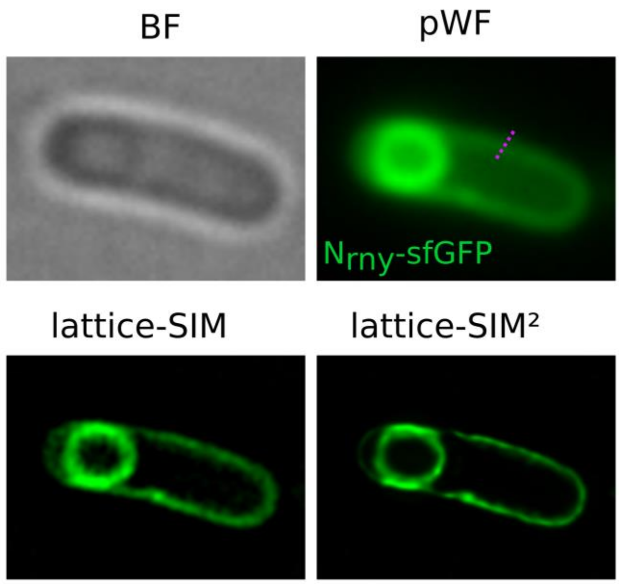


**C**

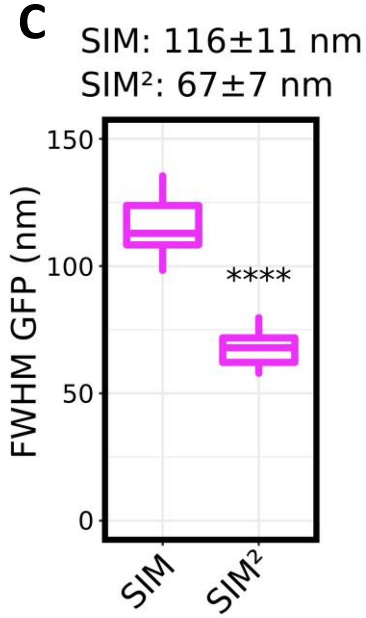

### Figure S5

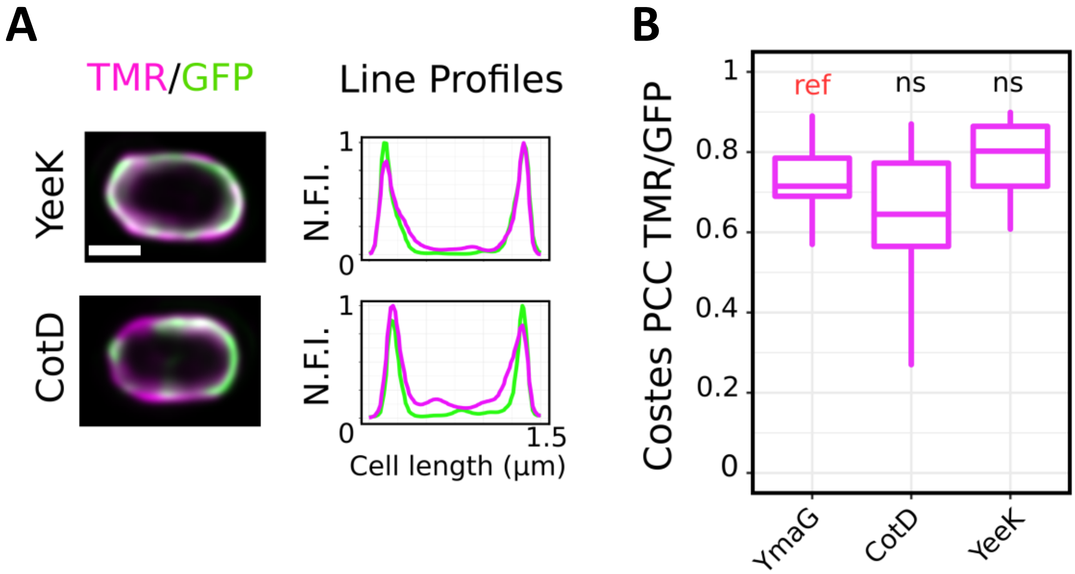

A
TMR/GFP
Line Profiles
YeeK
CotD
N.F.I.
1
0
1.5
Cell length (µm)
B
Costes PCC TMR/GFP
1
0.8
0.6
0.4
0.2
0
ref
ns
ns
YmaG
CotD
YeeK

Figure S6

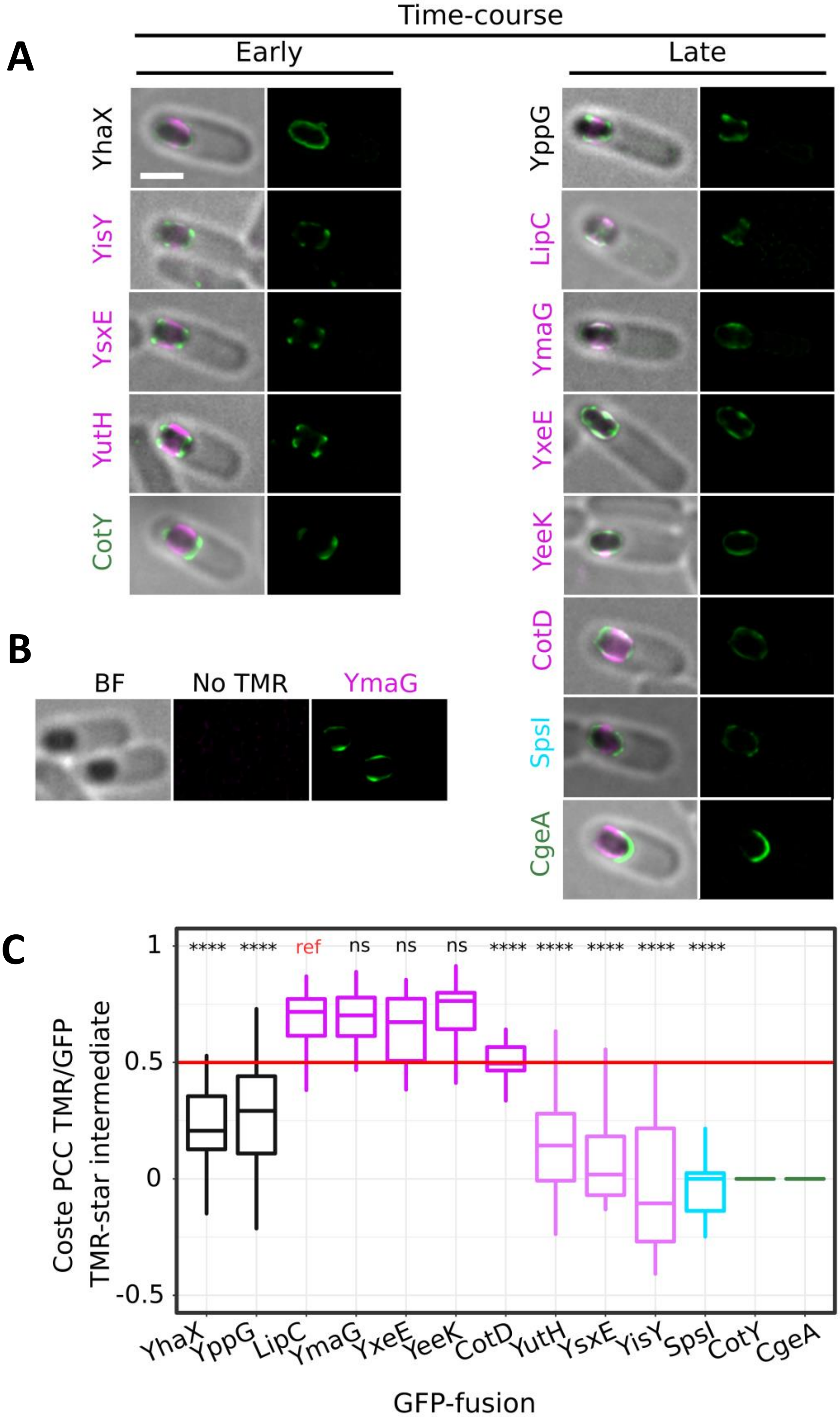
A
Time-course
Early
Late
YhaX
YisY
YsxE
YutH
CotY
YppG
LipC
YmaG
YxeE
YeeK
CotD
SpsI
CgeA
B
BF
No TMR
YmaG
C
Coste PCC TMR/GFP
TMR-star intermediate
1
0.5
0
-0.5
**** **** ref ns ns ns **** **** **** **** ****
YhaX
YppG
LipC
YmaG
YxeE
YeeK
CotD
YutH
YsxE
YisY
SpsI
CotY
CgeA
GFP-fusion

Figure S7

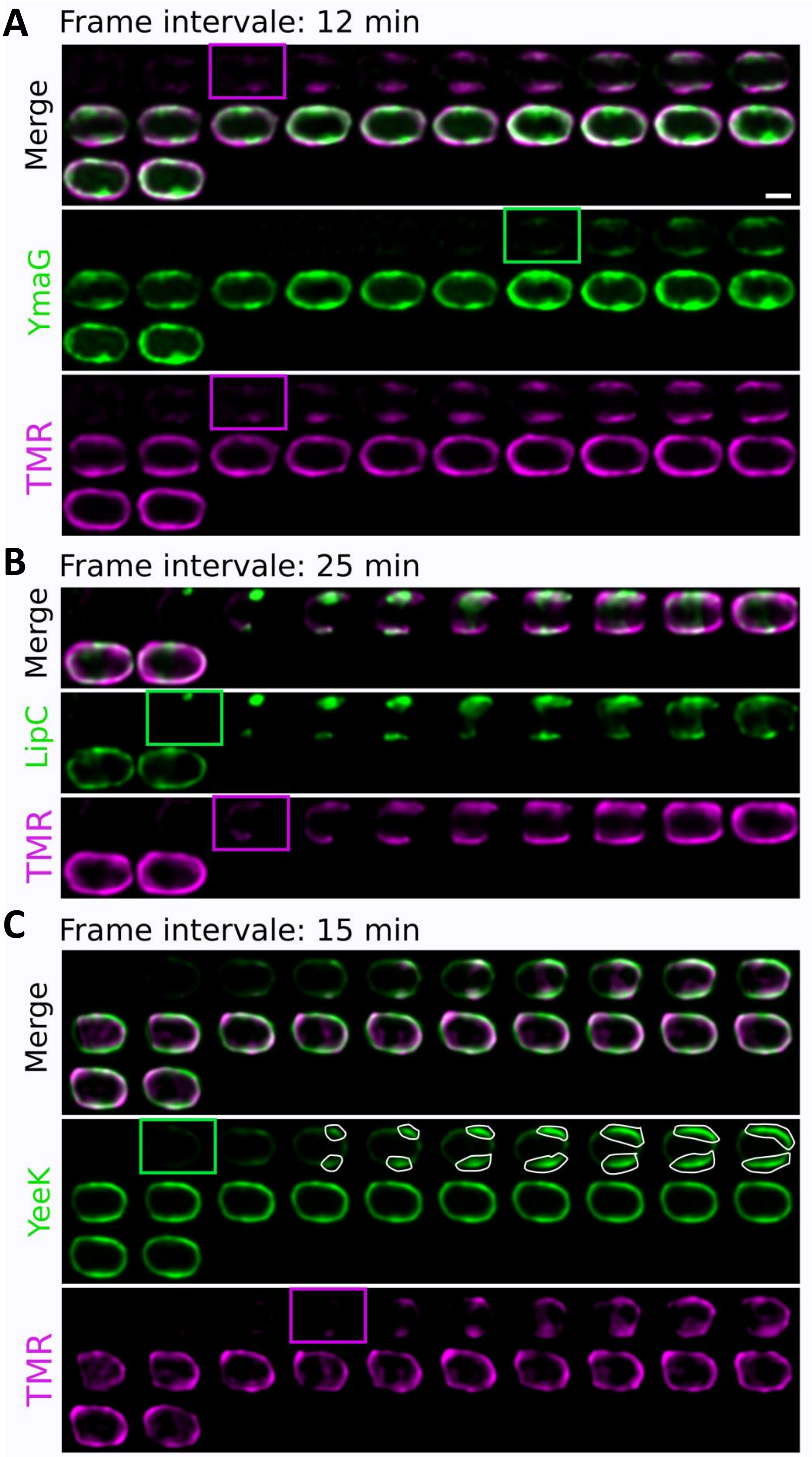
A
Frame intervale: 12 min
Merge
YmaG
TMR
B
Frame intervale: 25 min
Merge
LipC
TMR
C
Frame intervale: 15 min
Merge
YeeK
TMR

Figure S8

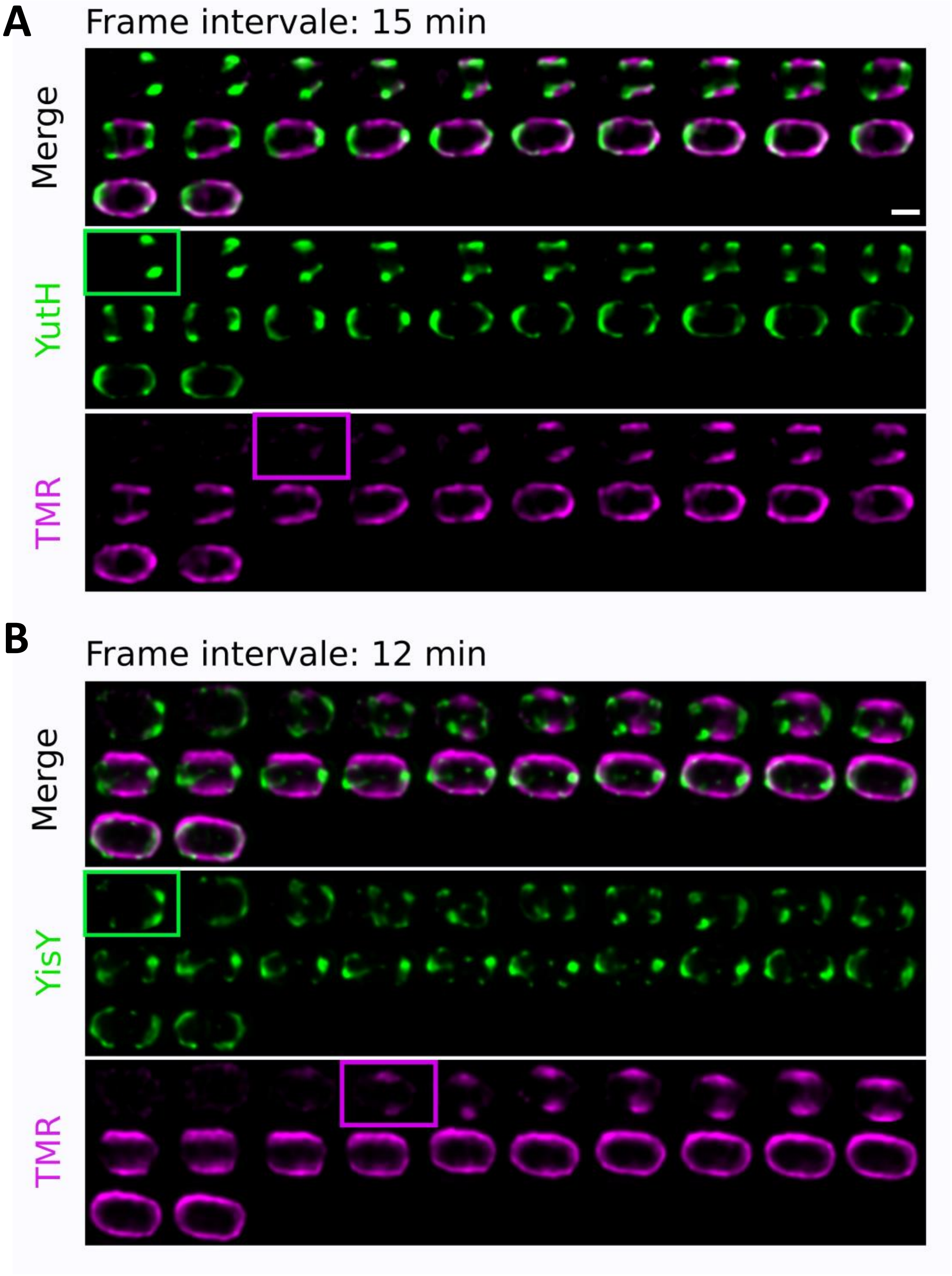
A
Frame intervale: 15 min
Merge
YutH
TMR
B
Frame intervale: 12 min
Merge
YisY
TMR

### Figure S9

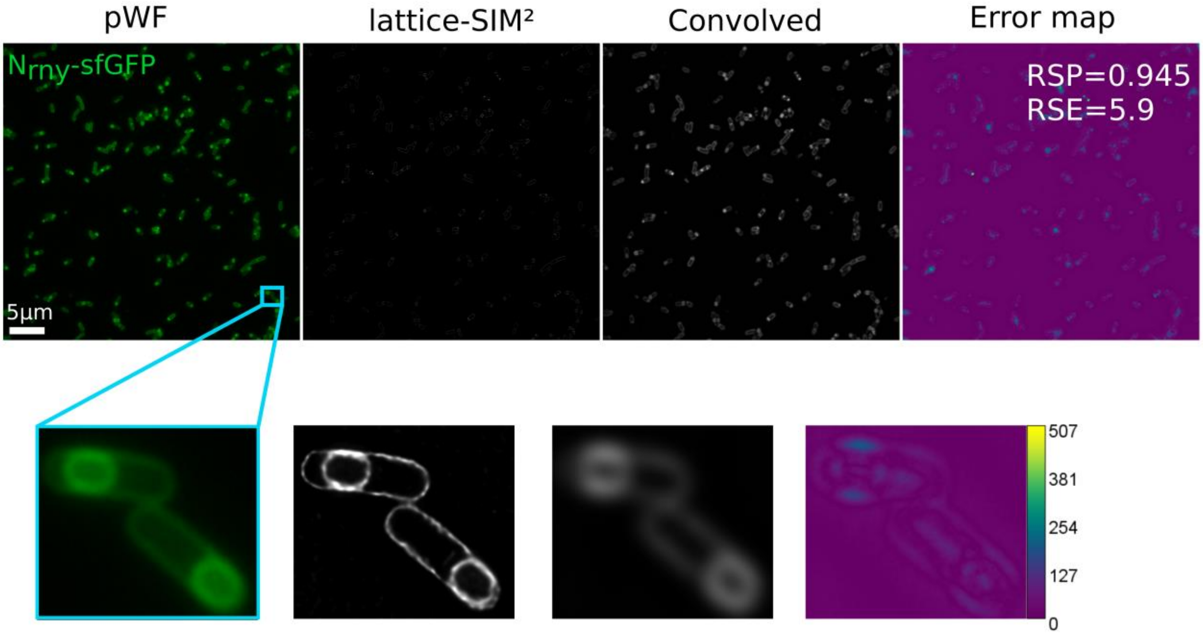
pWF
lattice-SIM²
Convolved
Error map
Nrny-sfGFP
RSP=0.945
RSE=5.9
5µm
507
381
254
127
0
